# Two-photon voltage imaging with rhodopsin-based sensors

**DOI:** 10.1101/2024.05.10.593541

**Authors:** Christiane Grimm, Ruth R. Sims, Dimitrii Tanese, Aysha S. Mohamed Lafirdeen, Chung Yuen Chan, Giulia Faini, Elena Putti, Filippo Del Bene, Eirini Papagiakoumou, Valentina Emiliani

**Affiliations:** Institut de la Vision, Sorbonne Université, INSERM, CNRS, F-75012 Paris, France

**Keywords:** two-photon microscopy, voltage indicators, microbial rhodopsin, scanless imaging, fast functional imaging, action potential detection

## Abstract

The recent advances in sophisticated optical techniques, coupled with two-photon sensitive genetic voltage indicators (GEVIs), have enabled in-depth voltage imaging *in vivo* at single spike and single-cell resolution. To date, these results have been only achieved using ASAP-type sensors, as the complex photocycle of rhodopsin-based voltage indicators posed challenges for their two-photon use, restricting their application to one-photon approaches. In this work, we demonstrate that rhodopsin-based GEVIs (FRET-opsin) can be used under two-photon illumination when their peculiar light intensity dependence of kinetics and sensitivity are considered. We rationally engineer a fully genetically-encoded, rhodopsin-based voltage indicator with the brightest known fluorophore AaFP1, Jarvis, and demonstrate its utility under both one- and two-photon illumination. We also showed two-photon usability of the similar FRET-opsin sensor pAce. Our comparison of 2P scanless with fast 2P scanning illumination revealed that the latter approach is less suitable for this class of indicators and, on the contrary, both sensors responded well when scanless approaches were used. Furthermore, utilising Jarvis, we demonstrated high-fidelity, high-SNR action potential detection at kilohertz-imaging rates both in mouse hippocampal slices and in zebrafish larvae. To the best of our knowledge, this study represents the first report of a fully genetically-encoded rhodopsin-based voltage indicator for high contrast action potential detection under two-photon illumination *in vitro* and *in vivo*.

## Introduction

Monitoring the activity of groups of neurons at single cell resolution is indispensable to understand how a complex brain computes. With the rise of genetically-encoded calcium indicators (GECIs), the invasive, low-throughput nature of electrical patch-clamp recordings could be overcome such that two-photon (2P) calcium imaging is now the gold-standard method for observing the activity of a group of neurons *in vivo*. Despite its popularity, calcium imaging has limitations like the inability to resolve subthreshold activity, fast trains of action potentials (APs), or hyperpolarisation, which could be overcome by directly monitoring the membrane potential instead of secondary calcium rise.

Hence, over the last decade genetically-encoded voltage indicators (GEVIs) were increasingly used to monitor neuronal activity of genetically defined cells, despite the technical challenges voltage imaging poses. In contrast to GECIs located in the cytosol, voltage indicators are restricted to the plasma membrane, which complicates protein engineering efforts and reduces the number of fluorophores available for imaging^1^. Together with the kilohertz acquisition rates necessary to resolve fast voltage changes like APs, this entails that optical recordings of membrane potentials operate on a tight photon budget. Consequently, despite the large number of GEVIs available to date^2^, there is an ongoing effort to improve existing sensors and develop new ones to enable *in-vivo* experiments at higher signal-to-noise ratios (SNR).

Existing GEVIs that were successfully used for the detection of APs, fall into two groups i) ASAP-type sensor using the voltage sensing domain of a phosphatase fused to a circularly permutated GFP^3^ and ii) rhodopsin-based sensors that are employing a microbial rhodopsin as voltage sensing domain either fused to a fluorescent protein called FRET-opsin sensors^4^ or using the fluorescence of the rhodopsin itself as output^5^. The group of rhodopsin-based sensors additionally comprises the chemigenetic indicators^6^, which do use a microbial rhodopsin as voltage sensing domain, but fused to a dye-capture protein domain that irreversibly binds a synthetic fluorophore. Through the use of the microbial rhodopsin they can still benefit from genetic targeting, but are not fully genetically-encoded as addition of the synthetic dye is needed.

To facilitate *in-vivo* AP detection, especially in depth or densely labelled preparations, the ideal GEVI should not only be bright and have fast kinetics, but should also be compatible with two-photon (2P) excitation. However, to date, only the ASAP-type sensors have been successfully used for AP detection under 2P illumination^7–12^. Although rhodopsin-based sensors are intrinsically brighter than the ASAP-type ones^6,13^, they are considered incompatible with 2P illumination and no 2P detection of APs has been reported to date^14,15^.

Here, we report the rational design of a fully genetically-encoded rhodopsin-based voltage indicator following the FRET-opsin design (Figure 1A) using the well-characterised *Acetabularia* rhodopsin (Ace)^16,17^. We replaced the fluorophore mNeonGreen with the brightest known fluorescent protein AaFP1^18^ and obtained a bright and fast indicator, which we call Jarvis, short for: **J**ust **a**nothe**r v**oltage **i**ndicating **s**ensor. We demonstrate that Jarvis can be used under 2P scanless illumination to detect action potentials in organotypic hippocampal slices and zebrafish larvae at kilohertz frame rates. To the best of our knowledge, this is the first report of a fully genetically-encoded FRET-opsin voltage indicator for action potential detection under 2P illumination. To understand why previous reports concluded a loss of voltage sensitivity under 2P illumination for this group of indicators, we performed a detailed comparison of parallel and scanning illumination for 2P voltage imaging with the rhodopsin-based sensors Jarvis and pAce and the ASAP-type sensor JEDI-2P. We conclude that ASAP-type sensors perform equally with both parallel and scanning illumination strategies, while positive-going FRET-opsin sensors showed a significantly higher SNR with low-irradiance scanless illumination approaches.

**Figure 1:**
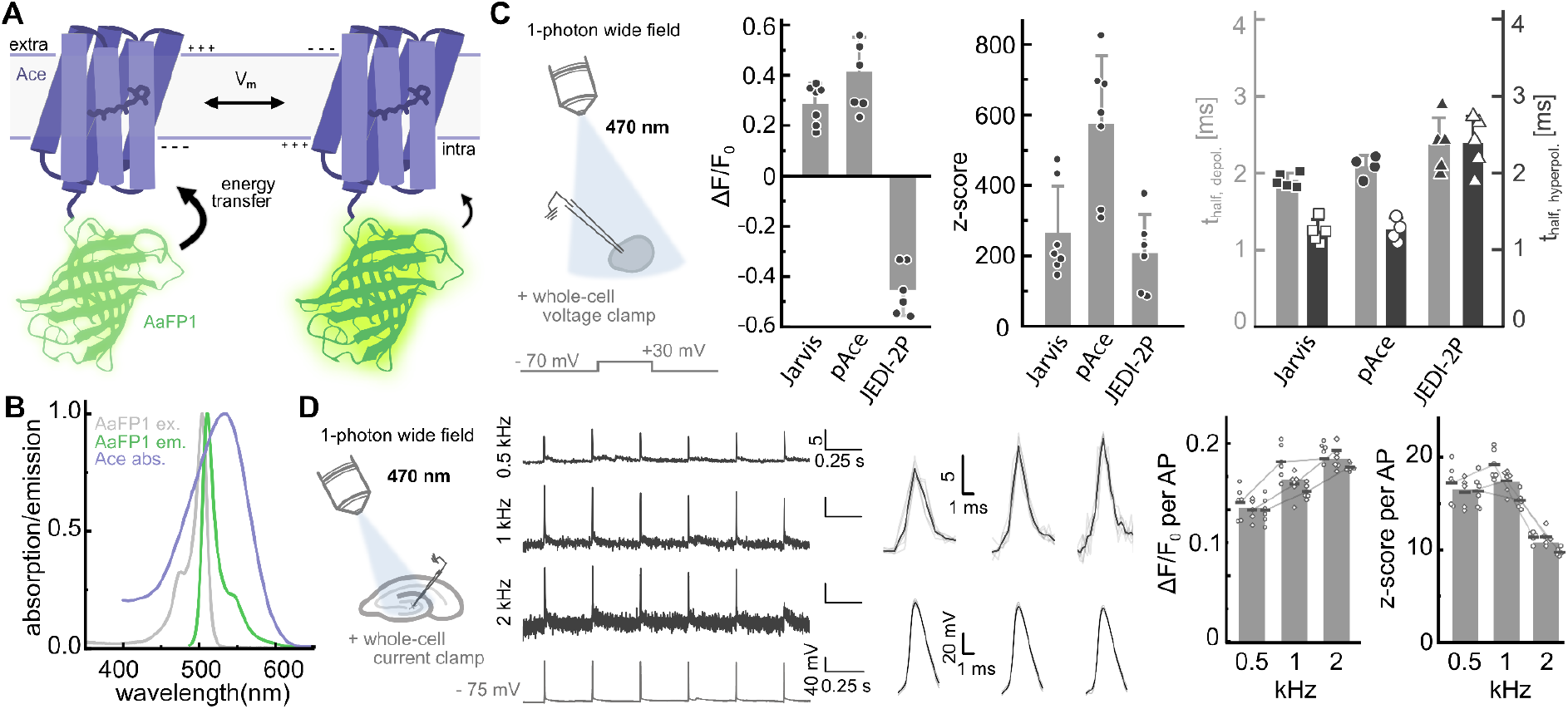
Engineering and one-photon characterisation of Jarvis. **A)** Schematic illustration of FRET-opsin voltage sensor with microbial rhodopsin in purple and fluorophore in green. The rhodopsin adopts two different states depending on the membrane potential leading to different efficiencies of the energy transfer from the fluorophore and hence brightness. **B)** Absorption spectrum of *Acetabularia* rhodopsin (purple) together with excitation (grey) and emission (green) spectrum of *Aa*FP1. **C)** *In-vitro* characterisation of Jarvis compared to pAce and JEDI-2P in cultured ND7/23 cells using 1P wide field illumination (470 nm LED, 10 mW/mm^2^) with camera detection (50 Hz) and simultaneous whole-cell voltage-clamp (−70 mV to 30 mV for 100 ms). ΔF/F_0_ and z-score extracted from imaging traces at the end of the voltage step while kinetics were extracted (monoexponential fit) from ΔF/F_0_ traces acquired at 1 kHz. **D)** 1P wide field imaging (470 nm LED, 16 mW/mm^2^) and simultaneous whole-cell current-clamp recordings in organotypic hippocampal slices sparsely expressing Jarvis after electroporation (room temperature). Per recording six APs were electrically injected and imaged at frame rates from 0.5 to 2 kHz with an CMOS camera. Left: Exemplary traces from one cell at all frame rates; electrophysiology trace from 1 kHz recording. Middle: Overlay of extracted APs for each frame rate respectively. Right: Summary of ΔF/F_0_ and z-score per AP; Points denote individual APs, horizontal lines average per cell, bars are average over all three cells.

## Results

### Engineering and one-photon characterisation of Jarvis

As the detection of action potentials demands kilohertz acquisition rates, 2P voltage imaging is operating on a low photon budget. To tackle this problem there is a continued need to develop brighter voltage sensors, which we approached by replacing the fluorophore in the well-established FRET-opsin design (Figure 1A) with the brightest known fluorophore *Aequoria australis* FP1 (*Aa*FP1)^18^. Besides the high brightness *Aa*FP1 displays a narrow excitation spectrum and an extraordinary quantum yield of 97%, which together with the near complete overlap of its emission spectrum with the absorption spectrum of the *Acetabularia* rhodopsin (Ace) (Figure 1B) should render it an excellent FRET donor to Ace.

The first variants were designed like VARNAM^19^ using the same cutting site for Ace and the same linker sequence. To decide on the fusion site for *Aa*FP1 we performed a structural alignment with mRuby3 in PyMOL^20^ to find the fusion site structurally similar to that in VARNAM. Ultimately, we cut seven amino acids from the unstructured N-terminus of *Aa*FP1 for the first fusions (Supplement figure S1), but neither PA007 nor PA008 expressed well in ND7/23 cells. Adding back the complete N-terminus of *Aa*FP1 resulted in well-expressed variants, which were then further characterised. PA004, PA012 and PA013 are negative-going variants with different membrane targeting and expression enhancement sequences, of which PA004 had the best performance (Supplement figure S1). PA018 includes the Positron mutations^21^ and showed a positive-going response, but only a 10% ΔF/F_0_ for a 100 mV step. Adding the Voltron2 mutation^22^ A165D onto PA013 did not result in an improvement of the sensitivity (data not shown). Finally, transferring the pAce mutations R121K, N124D, D135N, W221F^17^ and linker onto PA013 yielded PA020, a bright and well-expressed variant, which we called Jarvis, short for **J**ust **a**nothe**r v**oltage **i**ndicating **s**ensor.

We expressed Jarvis and the above described variants in ND7/23 cells and performed whole-cell voltage-clamp recordings, stepping the membrane voltage from −70 mV to 30 mV, to compare their 1P performance to pAce and the ASAP-type GEVI JEDI-2P^9^. Jarvis showed a comparable ΔF/F_0_ as pAce (0.29±0.08 *vs*. 0.41±0.14, p = 0.16) while JEDI-2P had the highest ΔF/F_0_ (0.46±0.10) in this experiment. Despite the lower ΔF/F_0_ both Jarvis (265±132) and pAce (575±193) showed a higher average z-score than JEDI-2P (208±110), which we attribute to the higher brightness of the FRET-opsin sensors. Comparing the time constants for the fluorescent changes upon depolarisation we found no significant difference between Jarvis, pAce and JEDI-2P (1.9±0.1 ms, 2.1±0.2 ms, 2.4±0.4 ms) while the hyperpolarising kinetics were slower in JEDI-2P (1.3±0.2, 1.3±0.2, 2.4±0.4 ms; Jarvis *vs*. JEDI-2P: p = 0.016, pAce vs. JEDI-2P: p = 0.008) (Figure 1C).

Lastly, for the 1P characterisation, we electroporated hippocampal organotypic slices to obtain sparse expression of Jarvis in neurons and performed patch-clamp recordings with simultaneous voltage imaging using 1P wide field illumination and camera detection with an CMOS camera. Electrically evoked APs were readily detectable in single trials at acquisition rates of up to 2 kHz (mean z-score per AP: 0.5 kHz 16.6±0.6; 1 kHz 17.3±1.9; 2 kHz 10.8±0.9). ΔF/F_0_ per AP was observed to rise as a function of acquisition rate from 0.20±0.01 at 0.5 kHz over 0.25±0.02 at 1 kHz to 0.28±0.01 at 2 kHz (Figure 1D).

#### Holographic two-photon voltage imaging with Jarvis in slices

Following the 1P characterisation of Jarvis, we generated a Jarvis-encoding adeno-associated virus (AAV2/9) to express Jarvis panneuronally in organotypic hippocampal slices. We employed two different strategies for virus delivery: bulk transduction and microinjections. As seen from the confocal images (Supplement figure S3), we found high expression levels 10 to 13 days after transduction in both cases, throughout the slice for the bulk transduction and localised around the injection site for the microinjection (*cf*. Supplement figure S3A and B).

We then performed whole-cell current clamp recordings and simultaneous 2P scanless imaging using targeted holographic illumination (Figure 2A). The optical setup to generate a temporally-focused (TF) computer-generated holographic (CGH) spot was as described previously^12^. Here, we used a high-repetition laser tuned to 940 nm to project a disk with a radius and axial extent of **∼**10 μm, approximately the size of the soma of the neurons in the polymorphic layer of dentate gyrus predominantly patched (Figure 2B). We recorded from Jarvis-expressing cells and first determined resting membrane potential, input resistance and rheobase, where we found no difference to non-expressing neurons (Supplement figure S4A). Subsequently, we performed current-clamp recordings and electrically injected APs at 10 Hz, 25 Hz and 50 Hz while simultaneously imaging the fluorescence change of the patched cell at a 1 kHz acquisition rate (Figure 2C). For each cell, we injected between 9 and 13 APs and analysed their features in the imaging and electrical traces (Supplement figure S4B). We extracted the ΔF/F_0_ and z-score per injected AP for each neuron and found an average ΔF/F_0_ per AP across all neurons of 0.24±0.06 and an average z-score of 12.3±2.7 per AP (Figure 2D). For a subset of cells we compared ΔF/F_0_ and z-score per AP at different acquisition rates of 0.5 kHz, 1 kHz, and 2 kHz on the same cell (Supplement Figure S5A). We found that both ΔF/F_0_ and z-score per detected AP on average decreased when the acquisition rate was lowered to 0.5 kHz. When we increased the acquisition rate to 2 kHz (at constant irradiance) ΔF/F_0_ further increased, but the z-score per AP plateaued (Supplement figure S5B). This suggests 1 kHz as the optimal acquisition rate for Jarvis.

**Figure 2:**
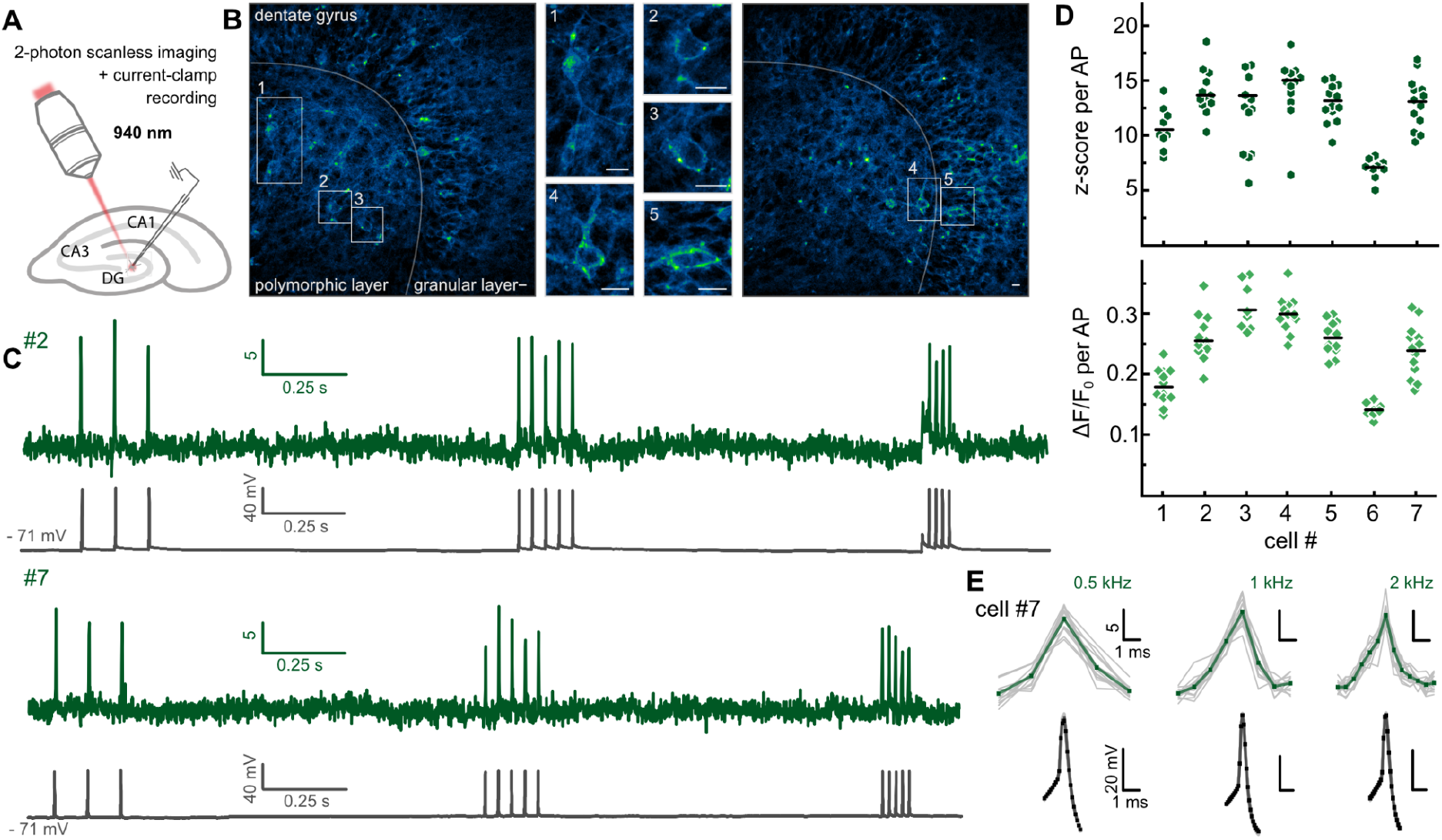
Holographic two-photon voltage imaging with Jarvis. **A)** Organotypic hippocampal slices virally transduced with Jarvis. Expressing neurons were patched and simultaneously imaged with an sCMOS camera, illuminated with a temporally-focused holographic spot (940 nm, **∼**20 μm diameter, **∼**10 μm axial extent, 0.66 mW/μm^2^) projected onto the soma of the neuron (33°C). **B)** 2P scanning (900 nm, 5-9 mW) images of dentate gyrus of fixed organotypic slices expressing Jarvis. Zoom-ins 1-5 show exemplary cells in the polymorphic layer that were predominantly patched. Scale bar always corresponds to 10 μm. **C)** Exemplary z-score trace (green) from imaging neuron #2 (1 kHz acquisition) with corresponding electrophysiology trace (grey). APs were evoked electrically throughout the 5 s long recording at 10 Hz, 25 Hz and 50 Hz. **D)** z-score and ΔF/F_0_ per successfully detected AP for all patched neurons displayed as individual data points together with the average (black line). **E)** Comparison of AP waveform at 0.5 kHz (left, neuron #4), 1 kHz (middle, neuron #3) and 2 kHz (right, neuron #5) acquisition rates with optical z-score (green) at top and corresponding electrical trace (grey) at bottom; bold line average and thin lines individual APs.

Taken together and contrasting previous reports^14,15^ we were able to demonstrate that rhodopsin-based GEVIs are usable with 2P illumination and allow reliable AP detection at similarly high SNR as reported for ASAP-type sensors^12^. Hence, 2P imaging is not an intrinsic limitation of FRET-opsin GEVIs. Nevertheless, we wondered whether this observed 2P sensitivity was a feature of our new sensor Jarvis or a consequence of our parallel, scanless illumination strategy. To answer this question, we performed a side by side comparison with pAce, as another rhodopsin-based sensor, and JEDI-2P in cultured cells. We patched expressing cells and stepped the holding potential from −70 mV to 30 mV for three times 3 ms while simultaneously imaging the fluorescence changes under parallel illumination at 940 nm (Supplement figure 6A). Both Jarvis and pAce clearly showed a voltage dependent fluorescence change for these AP-like changes of the membrane potential.

#### Comparison of two-photon scanning and parallel illumination

To assess the impact of the illumination strategy on the 2P response of the sensors, we designed an experiment to compare performance of Jarvis, pAce and JEDI-2P under conventional 2P scanning *versus* parallel, scanless illumination on the same cell (Figure 3A, Supplement figure S7A). In general, the two-photon signal S is proportional to the illuminated volume V containing the fluorophore, the duration of the illumination τ_dwell_ and the square of the irradiance P according to *S* ∝ *P*^2^ * *V* * τ_*dwell*_. We set up the experiment such that scanning and parallel illumination covered comparable areas (same cell) (Supplement figure S7) resulting in the irradiance P and dwell time τ_dwell_ being the predominantly different parameters between the two illumination approaches. While scanning takes advantage of increased irradiance in the diffraction limited spot, the beam has to be moved quickly to cover the whole area resulting in microsecond dwell times. Conversely for the parallel illumination, irradiances are two orders of magnitude lower, but the whole area is illuminated in parallel resulting in millisecond dwell times.

**Figure 3:**
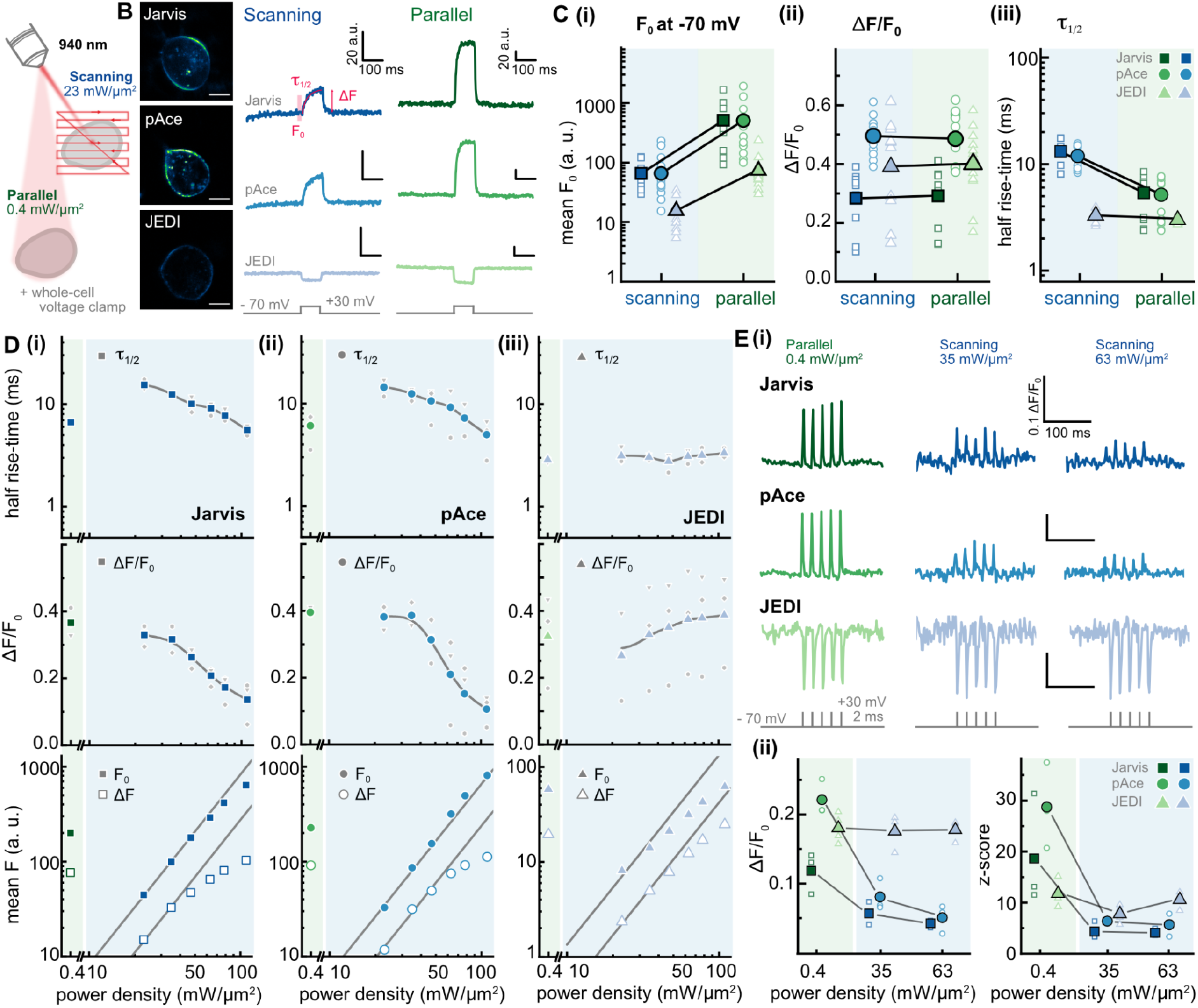
Comparison of scanning and parallel illumination for two-photon voltage imaging. ND7/23 cells were patched and the holding potential was stepped from −70 mV to +30 mV for 100 ms. **A)** The voltage response was imaged either scanning a frame-like trajectory over the cell (1.804 ms per trajectory, 4 μs pixel dwell time) or using parallel illumination of the whole cell with a large Gaussian spot (0.4 mW/μm^2^). Wavelength was 940 nm and acquisition done with a PMT for both illumination strategies (554 Hz). **B)** Images of cells expressing Jarvis, pAce or JEDI (scale 10 μm) and exemplary raw fluorescence traces for each sensor with scanning (blue) and parallel (green) illumination on the same cell, respectively. **C)** Average (i) baseline fluorescence F_0_ at −70 mV 250 ms after light on, (ii) relative fluorescence change ΔF/F_0_ during the voltage step and (iii) half-rise time of the fluorescence change for Jarvis (square), pAce (circle) and JEDI (triangle) during scanning (blue, 23 mW/μm^2^) or parallel illumination (green, 0.4 mW/μm^2^); filled symbols average and open symbols individual recordings. **D)** Half rise-time (top), ΔF/F_0_ (middle) and average F (bottom) at rising scanning powers from 23 mW/μm^2^ to 109 mW/μm^2^ (blue) and corresponding value from parallel illumination (green) for (i) Jarvis, (ii) pAce and (iii) JEDI expressing cells; n=3 each, whole titration on same cell. Top and middle panel show average values (colored) and individual data points (grey). Lower panels show average values for F_0_ and absolute fluorescence change ΔF; lines have a fixed slope of 2. **E)** (i) ΔF/F_0_ traces resulting from five short AP-like depolarisations imaged with parallel (green) and scanning (blue) illumination at two power densities for cells expressing Jarvis, pAce, or JEDI; traces were detrended to account for bleaching; comparison on the same cells respectively. (ii) Average ΔF/F_0_ and z-score for each cell obtained per depolarisation.

For an initial comparison, we chose a moderate irradiance of 23 mW/μm^2^ for scanning and a rather low irradiance of 0.4 mW/μm^2^ for the parallel illumination. We performed whole-cell voltage clamp recordings and stepped the membrane potential from −70 mV to 30 mV for 100 ms while simultaneously imaging the corresponding fluorescence change of the same cell either under scanning or parallel illumination (Figure 3B). We restricted this analysis to cells with similar ΔF/F_0_ between the two illumination methods; cells were excluded when ΔF/F_0_ was > 25% different between the two approaches. In these conditions, for all three constructs and all cells, the resulting baseline fluorescence F_0_ after 200 ms illumination at −70 mV was higher with parallel than scanning illumination (Figure 3C). Unexpectedly, we found that for the rhodopsin-based indicators Jarvis and pAce the fluorescence change due to the voltage step had a significantly slower rise time with scanning than with parallel illumination while for the ASAP-like indicator JEDI-2P there was no dependence on the illumination mode (Figure 3C (iii)). We speculated that this indicates light modulation of the voltage dependent transition for the rhodopsin-based indicators, suggesting that a variation of the illumination power could presumably affect the kinetics of this transition. Indeed, when we performed the same experiment at rising scanning powers from 23 mW/μm^2^ to 109 mW/μm^2^ (same cell), we found that for Jarvis and pAce the half-rise time is accelerated with increased irradiance while for JEDI-2P it was unchanged over the whole range (Figure 3D, top) confirming that for Jarvis and pAce the voltage dependent transition can be modulated by light.

In addition to the half-rise times, we extracted the ΔF/F_0_ from these recordings and found another effect dependent on the irradiance. For Jarvis and pAce the ΔF/F_0_ was significantly decreased with rising power densities dropping to less than half at the highest power densities, while for JEDI-2P it remained constant over the whole range (Figure 3D, middle). To further investigate this result, we separately plotted the F_0_ and the absolute ΔF for rising power densities (Figure 3D, bottom). We found that for Jarvis and pAce F_0_ was rising as expected for two-photon excitation following a power law with an exponent of two (with slight deviations due to bleaching), while the ΔF saturated at higher irradiances, which explains the observed reduction in ΔF/F_0_. For JEDI-2P, on the other hand, we did not observe any saturation of ΔF; here F_0_ and ΔF increased synchronously over the whole range of irradiances, maintaining constant ΔF/F_0_. We repeated the experiment using a CMOS camera instead of a PMT for detection and confirmed the results (Supplement figure S8A and B). Camera detection additionally allowed to not only target the illumination to the same cell, but also to average the fluorescence signal from exactly the same region of interest for scanning and parallel illumination. We again increased the irradiance for the scanning approach, here until it resulted in the same F_0_ as the parallel illumination of this cell. At similar F_0_ for scanning and parallel illumination JEDI-2P showed the same ΔF while Jarvis and pAce-despite equal F_0_-showed a significantly lower ΔF/F_0_ with scanning illumination (Supplement figure S8C).

We revealed that with scanning illumination at low and medium irradiances the fluorescence change rises slowly and that this rise time can be accelerated by incrementing the irradiance, but at high irradiances the amplitude of the ΔF/F_0_ reduced significantly. Hence, we were interested how the combination of these two effects affects the detection of shorter AP-like changes of the membrane potential. To examine this, we stepped the holding potential from −70 mV to 30 mV for five times 2 ms and compared the results for parallel (0.4 mW/μm^2^) and scanning illumination (35 and 63 mW/μm^2^) on the same cell expressing Jarvis, pAce or JEDI-2P, respectively (Figure 3E (i)). For JEDI-2P we found no difference in the ΔF/F_0_ between all illumination conditions, while for Jarvis and pAce the ΔF/F_0_ per depolarisation was significantly reduced with scanning illumination and further reduced at the higher irradiance. This was in line with the previous experiments and demonstrates that spike detection with rhodopsin-based indicators resulted in low SNRs with a scanning approach and was significantly improved with parallel illumination.

### Action potential detection in zebrafish larvae

Lastly, we wanted to show that rhodopsin-based sensors can be used to detect APs under scanless 2P illumination *in vivo*. We transiently expressed Jarvis in zebrafish larvae and performed imaging experiments 4-6 days post fertilisation (dpf) on spontaneously active neurons in the olfactory bulb and adjacent telencephalon (Figure 4A and B). During the experimental session, we initially identified spontaneously active neurons under 1P wide field illumination (470 nm LED, 7 mW/mm^2^) and subsequently performed scanless 2P imaging on the same neurons (Supplement figure S9). We used a high repetition rate laser (80 MHz) tuned to 920 nm to create a temporally-focused low-NA Gaussian spot (0.61-0.76 mW/μm^2^) targeted to the soma of the neurons, while acquisition was done using a sCMOS camera at an effective frame rate of 991 Hz. We detected APs under 2P illumination in all imaged cells and readily identified neurons with different firing patterns (Figure 4C). The z-score per AP was large across neurons (Figure 4D) with an overall average of 7.7 ± 1.9 and statistically indistinguishable from 1P illumination (Supplement figure S9). We performed continuous recordings of up to 30 s under scanless 2P illumination for several cells with AP detection still possible after extended illumination (Figure 4E, Supplement figure S10).

**Figure 4:**
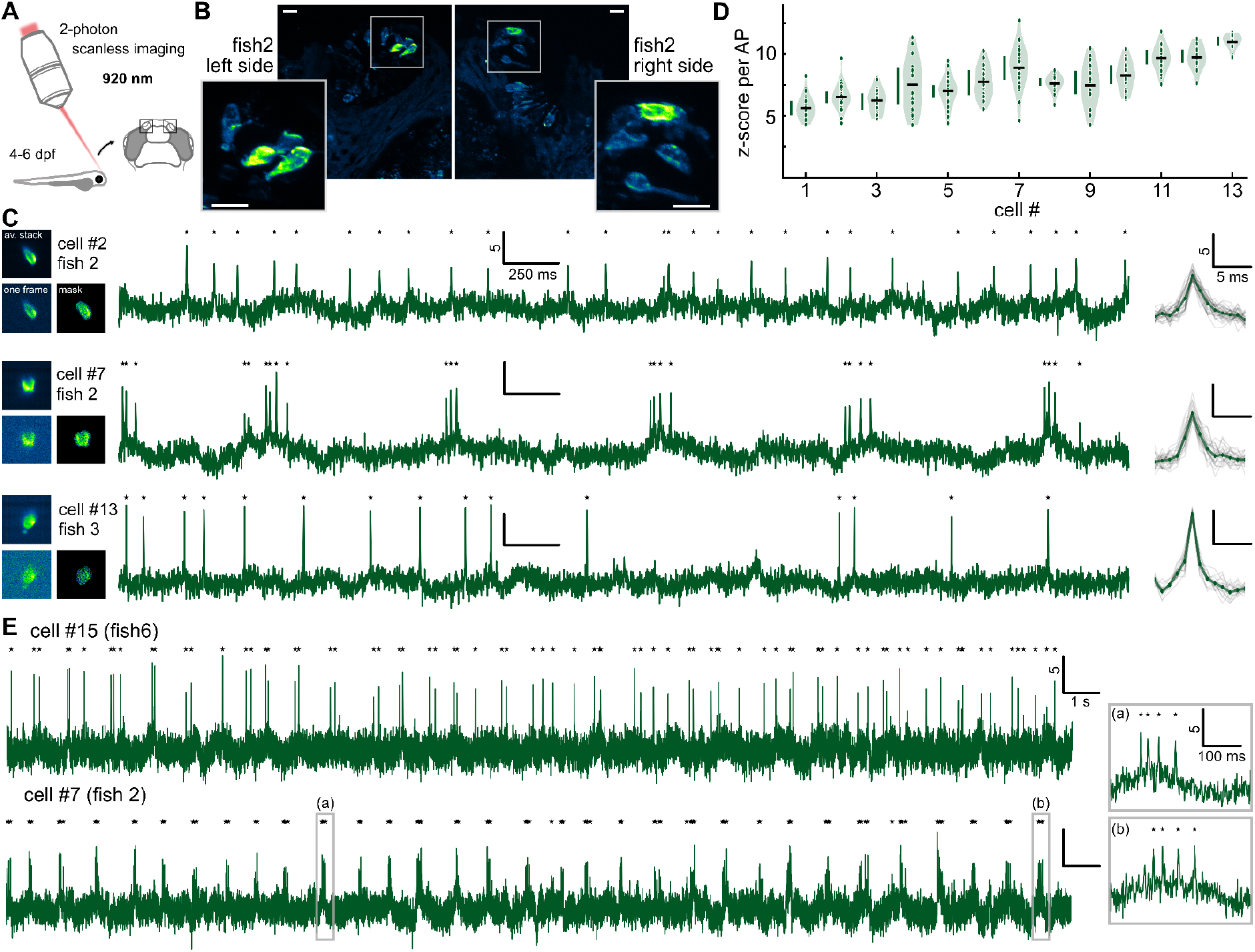
Scanless two-photon voltage imaging in zebrafish larvae using Jarvis. **A)** Parallel illumination was realised projecting a temporally-focused low-NA Gaussian spot (920 nm, 0.61-0.76 mW/μm^2^) onto the soma of olfactory sensory neurons. **B)** 2P scanning z-stack of Jarvis expressing neurons in an exemplary fish. Projection of several planes (2 μm) taken around the neurons. Scale bar corresponds to 10 μm. **C)** Representative z-score traces of three neurons from two different fish acquired at 991 Hz. Stars indicate detected APs with an overlay of all APs per trace on the right. Images depict the averaged stack, a single frame of the stack and the weights mask that was used for analysis. **D)** Summary violin plots of z-score per AP for 13 neurons from five fish recorded for five seconds. Points correspond to APs, black lines represent average per cell and green bars 25th and 75th percentile. **E)** Exemplary z-score traces from 30 s acquisitions (991 Hz) of two neurons. Insets show a zoom into a burst at the beginning (a) and end (b) of the recording.

## Discussion

Here, we demonstrate the rational engineering of a FRET-opsin voltage indicator, Jarvis, using today’s brightest known fluorophore *Aa*FP1. We expressed Jarvis in organotypic slices of mouse hippocampus and zebrafish larvae and successfully imaged APs under 2P illumination at kilohertz frame rates maintaining high SNR at acquisition speeds of routinely 1 kHz and even up to 2 kHz. To the best of our knowledge, this work represents the first demonstration of 2P voltage imaging with genetically-encoded FRET-opsin indicators for AP detection at high contrast *in vitro* and *in vivo* (SNR per AP: 7.7±1.9 at 1 kHz, Figure 4D).

In cultured cells under 1P illumination Jarvis performed similar to its parental sensor pAce with statistically insignificant, slightly lower ΔF/F_0_, but it had a lower SNR than pAce in cultured cells. After expression in neurons and under scanless 2P illumination, however, we obtained a higher z-score per AP for Jarvis than for pAce at the same imaging conditions (Supplement figure S11). On the one hand, the slightly lower ΔF/F_0_ shows that Jarvis can still be improved and serve as an excellent template for a high-throughput screen in the future. On the other hand, varying SNR between expression systems shows that voltage indicators are largely influenced by expression levels in the target cell type. Ultimately, performances are not only dependent on voltage sensitivity but also and especially on expression levels and membrane localisation, affecting overall brightness and signal-to-background ratio and therefore impacting SNR. Finally, our results exemplify that despite the modularity of the sensor design each combination warrants new optimisation, and not only changes in the voltage sensing domain or the linker, but even swapping the fluorophore can affect the performance of the sensor.

While Jarvis and pAce are similar, there is a distinct difference in the width of the excitation and emission spectra of the fluorophores mNeonGreen and *Aa*FP1^18^ (FWHM: 22 *vs*. 17 nm) (Supplement figure S11A), which has implications for the use of both indicators. As the spectrum of Jarvis is narrower we recorded significantly less crosstalk into the red PMT than with pAce (Supplement figure S11B). This feature suggests that spectral multiplexing with two voltage sensors as published recently^17^ would enable better unmixing with Jarvis. On the other hand, the broader spectrum of pAce entails that it can be more efficiently excited over a broader spectral range out of the peak excitation (Supplement figure S11C, D (i)), rendering pAce more compatible with lasers operating at wavelengths longer than 1000 nm (Supplement figure S11C, D).

Our comparison of scanning and parallel 2P illumination demonstrated that positive-going FRET-opsin voltage indicators can be used for 2P voltage imaging more efficiently when scanless approaches are employed that operate at lower irradiances than conventional 2P scanning approaches. This comparison revealed that the voltage sensitivity ΔF/F_0_ of rhodopsin-based indicators is a function of the irradiance and was decreasing for our investigated positive-going variants with rising power densities. This was an effect that we observed in all illumination modalities with evident reductions of ΔF for irradiances in the ranges: 1P wide field > 15 mW/mm^2^ (Supplement figure S12), 2P scanless > 0.8 mW/μm^2^ (Supplement figure S6) and 2P scanning illumination > 35 mW/μm^2^ (Figure 3). While 1P wide field and 2P scanless illumination approaches typically use power densities below these values, fast 2P scanning approaches^7,9,10,23,24^ easily operate in the regimes where the reduction of ΔF/F_0_ can be larger and hence reduce SNR (*cf*. Figure 3E). As the majority of 2P voltage imaging studies in the past relied on scanning approaches, we presume this to be one possible reason why the voltage sensitivity of rhodopsin-based sensors has not been reported before.

Additionally, from our comparison of scanning and parallel 2P illumination approaches, we conclude that FRET-opsin and ASAP-type sensors function in fundamentally different ways that have implications for their use. While for ASAP-type sensors ΔF was always a true fractional change of F_0_, those two parameters were more independent of each other for Jarvis and pAce, where F_0_ rose with the irradiance as expected, but ΔF saturated at high irradiances effectively reducing ΔF/F_0_. A comparable dependence of ΔF/F_0_ on power density was described for negative-going rhodopsin-only voltage indicators before, but here ΔF/F_0_ increased with rising irradiance^25^. For practical use of the indicators, this implies that for ASAP-type sensors approaches can focus on maximising F_0_, as the main determining factor for the expected ΔF/F_0_, to optimise SNR for AP detection. For pAce and Jarvis a high F_0_ was not a reliable indication for a high SNR; indeed our data suggests experiments to work best at the lowest possible irradiance that raises the signal sufficiently over the background.

For FRET-opsin sensors the efficiency of the energy transfer from the fluorophore to the microbial rhodopsin depends on the state of the microbial rhodopsin, which changes as a function of the membrane potential and provides the foundation for the optical read out^4,16^. For positive-going indicators like Jarvis and pAce energy transfer is efficient at more negative membrane potentials and less efficient at positive potentials (Figure 1A) resulting in an increase in fluorescence for depolarisations like APs. In our experiments the speed of the transition between the less and more fluorescent state evoked through a depolarisation from −70 mV to +30 mV was dependent on the illumination mode and faster with parallel illumination than with scanning illumination (see Figure 3). However, for scanning illumination the same kinetics could be achieved at increased irradiances supporting the conclusion of light-modulation of a transition. Mechanistically, these results could either be explained by a direct light-modulation of the voltage dependent transition of the microbial rhodopsin or by a light-activated transition that is necessary to populate the voltage dependent equilibrium as suggested previously for rhodopsin only indicators^25,26^.

Through all our experiments Jarvis and pAce consistently showed higher resting brightness F_0_ than the ASAP-type indicator JEDI-2P underlining the potential of FRET-opsin indicators for *in-vivo* voltage imaging approaches which inherently operate in a photon-limited regime. Moreover, rhodopsin-based sensors feature more linear sensitivity curves^4,16,17^ than the latest ASAP-type sensors^9,27^ (Supplement figure S2). This potentially allows higher responses to subthreshold voltage changes around neuronal resting potentials and higher versatility amongst different cell types, as for the same voltage change, comparable responses over a wider range of resting potentials can be expected.

In conclusion, our study confirms, consistent with previous studies^12^, that scanless 2P illumination is a viable strategy for high-contrast 2P voltage imaging to detect APs at high SNR *in vitro* and *in vivo*. We show that the rhodopsin-based voltage indicators Jarvis and pAce are highly compatible with 2P scanless illumination approaches. Hence, we clearly demonstrate for the first time that 2P scanless illumination approaches render a whole group of bright and sensitive voltage indicators functional under 2P illumination for AP detection *in vivo*. We gained viable insights on mechanistic aspects of positive-going, rhodopsin-based indicators and exposed differences to previously used ASAP-type indicators, which revealed beneficial experimental conditions to optimise SNR. Taken together, our study provides a new voltage indicator-Jarvis-together with an optimised illumination strategy and a set of tailored experimental parameters, which together lay a solid foundation for future voltage imaging approaches *in vivo* that necessitate 2P illumination, providing a viable strategy to image multiple cells in depth preserving high contrast.

## Material and Methods

### Molecular biology and Cell culture

Plasmids containing the coding sequences of all constructs were designed *in silico* using the SnapGene software (Dotmatics, Boston, USA) and synthesised/cloned into the same backbone (Supplement figure 1) under a CAG promoter by VectorBuilder (Chicago, USA). Site-directed mutagenesis was either provided by VectorBuilder or performed similar to the Quik-Change Site-Directed mutagenesis kit from Agilent Technologies (Santa Clara, USA) with the PrimeSTAR® GXL polymerase (TakaraBio, Shiga, Japan) and One Shot™ TOP chemically competent cells (Thermo Fischer Scientific, Waltham, USA). Plasmid amplification was either provided by VectorBuilder or performed with the Monarch® Plasmid miniprep kit from New England Biolabs (Ipswich, USA) or the NuceloBond Xtra Maxi kit with the NuceloBond Finalizer from Marchery-Nagel (Düren, Germany).

ND7/23 cells (Merck, Rahway, USA) were cultured in Dulbecco’s Modified Eagle’s Medium (high glucose) with 5% (v/v) foetal bovine serum and 1 μg/ml penicillin/streptomycin. For experiments cells were plated onto poly-D-lysine coated glass coverslips supplemented with 1 μM all*-trans* retinal and transfected using FuGENE® HD (Promega, Madison, USA) the next day. Experiments were carried out 36-45 hours after transfection.

### Organotypic slices culture

Slicing was performed according to the french law and guidelines of directive 2010/63/EU and the institutional guidelines on the care and use of laboratory animals. Organotypic slices of the hippocampus were prepared from P6-P8 C57BL/6J mouse pups as described before^28^ and transferred onto small pieces of sterilised polytetrafluorethylene membrane on membrane inserts (Millicell PICM0RG50). Slices were cultured at 37°C and at 5% CO_2_ in culture medium composed of 80% minimal essential medium and 20% inactivated horse serum supplemented with 1 mM L-glutamine, 0.1 mg/ml insulin, 0.00125% ascorbic acid, 13 mM D-glucose, 14.5 mM NaCl, 2 mM MgSO_4_,1.44 mM CaCl_2_ and 1 μg/ml penicillin/streptomycin; half of the medium was exchanged every 2-3 days.

Bulk electroporations were performed^29^ at 3 days *in vitro* with endotoxin-free plasmid preparations (> 1 μg/μl) of the plasmids used for transfecting ND7/23 cells, expressing pAce and Jarvis under a CAG promoter; experiments were performed 8-12 days after electroporation.

For adeno-associated virus (AAV) production, constructs were subcloned into a pAAV backbone with a synapsin promoter and AAV2/9 particles were produced in house at the Institut de la Vision. For viral expression, slices (DIV3 or 4) were either bulk transduced with 1 μl of diluted virus (1:5 to 1:10 from a titer of 4*10^14^ vg/ml for Jarvis and 1:10 from a titer of 5.3*10^14^ for pAce) or microinjected (same dilution) with a custom setup built around a picospritzer as described before^30^. Experiments were performed ten to 13 days after transduction.

### Preparation of zebrafish larvae

All procedures involving zebrafish were in accordance with national and European (2010/63/EU) guidelines and were approved by the authors’ institutional review boards and national authorities (French Ministry of Research, protocol ID: APAFIS# EU 1043-21323). Fish were maintained at 28.5 °C on a 14 h light/10 h dark cycle. Animals were housed in the animal facilities of the Vision Institute, which were built according to the local animal welfare standards. For the experiments we generated a transgenic fish line (*Xla*.*Tubb2-hsp70-ubc:kalt4*) by injecting the *Xla*.*Tubb2-hsp70-ubc:kalt4* plasmid with tol2 transposase mRNA at 25ng/μL at one-cell stage. To generate the plasmid, the *Xla*.*Tubb2* promoter sequence was obtained from a previously published plasmid gifted by M. Halpern^31^ The 3’ 638bp of the zebrafish *hsp70* promoter^32^ and the ubc intronic sequence^33^ were used to enhance the expression of the optimised Gal4, KalT4^34^. The four separate fragments were amplified and inserted into a vector containing the Tol2 sites^35^ using the Gibson Assembly Cloning Kit (New England Biolabs).

Experiments were performed on the above described line transiently expressing Jarvis (*Tg(Xla*.*Tubb2-hsp70-ubc:kalt4*). For that, Jarvis was subcloned using the Gibson Assembly Cloning Kit into a plasmid for expression in zebrafish under the 14UAS promoter (*Tg(14UAS:Jarvis*), by co-injecting the DNA construct (25 ng/uL) with tol1 transposase mRNA (25 ng/uL) in one-cell stage zebrafish embryos.

### Electrophysiology recordings

Patch pipettes were pulled from thick-walled, fire-polished borosilicate glass capillaries with filament (World Precision Instruments, Sarasota, USA) using a P-1000 micropipette puller (Sutter Instruments, Novato, USA) with a box filament. Pipette resistances ranged between 2-4 MOhm for recordings in ND7/23 cells and 3-6 MOhm for recordings in organotypic slices. Signals were amplified (MultiClamp 700B, Molecular Devices), filtered at 2/6/10 kHz and digitised at 10/50/500 kHz (DigiData 1400 or 1550B, Molecular Devices) using the Clampex software (Version 10.6 and 11.1).

In ND7/23 cells whole-cell voltage-clamp recordings were carried out at room temperature with an intracellular solution composed of 110 mM NaCl, 1 mM KCl, 2 mM CaCl_2_, 1 MgCl_2_, 1 mM CsCl and 10 mM HEPES (290 mOsm, pH 7.2) and extracellular solution: 140 mM NaCl, 2.5 mM KCl, 2 mM CaCl_2_, 1 mM MgCl_2_, 12.5 mM D-Glucose, 10 mM HEPES (310 mOsm, pH 7.2). The liquid junction potential between these solutions was calculated to be +1.2 mV and was not corrected. Pipette capacitance was compensated in cell-attached configuration before break-in. Series resistance was not compensated, but all protocols included an initial 10 mV depolarization to determine and monitor the quality of the patch, which was used to extract membrane (R_m_) and access resistance (R_a_)^36^. R_m_ was usually in the GOhm range (never below 400 MOhm) while R_a_ was below 15 MOhm (never above 25 MOhm). Additionally, for recordings with 100 mV depolarizing steps, the actual voltage change at the membrane (ΔV_m_) was estimated according to ΔV_m_ = (ΔV_command_*R_m_)/(R_m_+R_a_) and recordings discarded if the calculated ΔV_m_ was below 95 mV or changing more than 2 mV during the recording.

In organotypic slices whole-cell current-clamp recordings were performed at 33°C bath temperature (2P imaging) or at room temperature (1P imaging) as indicated in the respective figure. The intracellular solution was composed of 135 mM K-gluconate, 4 mM KCl, 10 mM HEPES, 10 mM Na-phosphocreatin, 0.3 mM Na_2_-ATP, 4 mM Mg-ATP and extracellular solution contained 125 mM NaCl, 2.4 mM KCl, 4 mM CaCl_2_, 4 mM MgCl_2_ (1.5 mM CaCl_2_, 1 mM MgCl_2_ for electroporated slices/1P experiments), 26 mM NaHCO_3_, 1.25 mM NaH_2_PO_4_, 25 mM D-Glucose, 0.5 mM ascorbic acid, which was bubbled with carbogen (95% O_2_ and 5% CO_2_) and continuously perfused the chamber at 1-2 ml/min. The liquid junction potential between these solutions was calculated according to the stationary Nernst-Planck equation^37^ using LJPcalc software to be +15.8 mV (15.2 mV for low Ca^2+^/Mg^2+^) and was corrected during analysis. After establishing whole-cell configuration passive membrane parameters and quality of the patch were assessed in voltage clamp mode by inducing a 10 mV, 100 ms depolarization; median R_m_ = 258±98 MOhm (always > 130 MOhm) and median R_a_ = 12±5 MOhm (always < 25 MOhm) across patched neurons. After switching to current-clamp mode, bridge balance capacitance was compensated with the internal circuits of the amplifier and neuronal parameters like resting potential, input resistance and rheobase were recorded right after break-in (Supplement figure S4). During the recordings, holding currents (−70 pA to 70 pA) were injected to keep the neurons at comparable membrane potentials between −75 mV and −80 mV.

Analysis of the electrophysiological data was performed with the Clampfit Software (version 11.1.0.23) and custom python (version 3.7.16) scripts built on the pyabf package (version 2.2.8).

### Confocal Imaging

For confocal imaging, organotypic hippocampal slices (prepared as described above) were individually transferred to a 24-well plate 10-12 days after transduction and fixed by submerging them in freshly prepared 4% paraformaldehyde for at least 30 minutes. After washing them with phosphate buffered saline thrice, they were mounted onto glass slides using Fluoromount-G mounting medium (Southern Biotech, Birmingham, AL, USA), covered with a coverslip, dried at room temperature, sealed with nail polish, and stored at 4°C in the dark.

Confocal images were acquired with a FluoView1000 confocal laser scanning microscope (Olympus, Tokyo, Japan) with a 20x, NA 0.85 oil immersion objective. Jarvis fluorescence was excited using a 488 nm argon laser using intensities between 2-4%. To image the whole slices 16 or 20 tiles (0.6 μm z-plane, 35-37 planes) were acquired, stitched together using the Fluoview software, exported as a tiff-file. Displayed images are a maximum intensity projection prepared with the Fiji software. For the acquisitions at higher zooms z-planes were 2 μm thick and three to four planes were average projected to obtain the displayed images.

### One-photon wide-field voltage imaging

The system (Supplement figure S7A) was custom-built around a commercial upright microscope (Axio Examiner, Zeiss). For 1P illumination a CoolLED (pE-4000, CoolLED) was coupled to the epifluorescence port of the microscope via a liquid light guide and the pE-Universal Collimator (CoolLED). Illumination duration and intensity were set with the analog outputs of the digitiser controlled by the Clampex software, no mechanical shutter was used. For all 1P experiments the 470 nm LED was used together with a filter block containing a 466/40 nm excitation filter, a 495 nm dichroic mirror and a 525/50 nm emission filter. The average power density out of the 20x objective (Zeiss, WPlan-Apochromat 20×/1.0 DIC) was 10 mW/mm^2^ if not stated otherwise. Acquisition was done with a DaVinci2K CMOS camera (SciMeasure) at 50 Hz or 1000 Hz (as indicated) without any binning. To conduct patch-clamp recordings, the microscope is equipped with a micromanipulator (Junior XR, Luigs and Neuman) and signals were amplified and digitised as described above.

Analysis of the 1P data is described below with the exception of Supplement figure S12, where we did not use a spatiotemporal correlation and used the same (binary) segmentation mask for all irradiances.

### Analysis of imaging data

If not stated otherwise in the respective method section, imaging datasets were exported as tiff files and analysed using a pipeline written entirely in Python, described previously^12^ and derived from^38^.

Briefly, a spatiotemporal correlation (8×8 pixel neighbourhood) was calculated on each raw dataset. The resulting 2D image was segmented using the random_walker function in scikit-image (beta: 130, mode: ‘cg’). This segmentation was then dilated using a disk radius 1.2 times the average cell diameter. The inverse of this mask was used to estimate the background. An initial fluorescence trace (mean pixel counts as a function of time) was calculated using the average value of each acquired frame, masked by the binary segmentation. A histogram of fluorescence counts was computed. The average “light-on” and “light-off” counts were estimated as the locations of the two largest peaks in this histogram. A condition trace corresponding to where the number of counts was greater than 3 standard deviations from the mean number of “light-off” counts was generated. Peaks in the derivative of this trace were used to identify timepoints where the light source was switched on and off and ultimately to calculate the indexes of “light-on” and “light-off” timepoints. Dark frames were calculated as the average of data of all “light-off” timepoints and subtracted from the raw dataset. Subsequent analysis was only performed for the dark-frame subtracted “light-on” timepoints. In cases where the light was switched on and off multiple times during a single acquisition (for instance the sensitivity curve datasets), each “light-on” period is hereafter termed an epoch. Next, the indexes corresponding to baseline and fluorescence transients were estimated. If the dataset consisted of multiple epochs, these indexes were estimated using the average epoch. For long voltage steps, these baseline and transient indexes were identified using the histogram approach outlined above. For action potential data, a z-score trace ((counts–mean(counts))/ (std_dev(counts)) was calculated and peaks with a z-score greater than 3 identified. The average width of these peaks was estimated, and indexes not within this boundary taken as baseline indexes. Fluorescence trends were estimated using an adaptive b-spline following^39^, built in scipy function: make_lsq_spline (knots: every 10 ms for the baseline indexes, order: 2). Trends were computed on a pixel wise basis and subtracted from the raw dataset. Clean fluorescence traces were then generated as follows. For long voltage steps the baseline fluorescence at each timepoint was replaced by the mean value of all time points and the transient fluorescence replaced by the mean transient fluorescence during each epoch. For data containing action potentials, an action potential template was calculated as the mean of all action potentials identified previously. A clean trace of zeros was generated and putative action potential indexes set to one. This clean trace was then convolved with the action potential template. A spatial filter was then generated by ridge regression (from sklearn.linear_model.Ridge) of the whitened clean trace against the whitened, detrended, segmentation masked, dataset. Final fluorescence traces were then calculated as the weighted spatial average of segmented pixels, with the background trace subtracted.

Two of the scanless 2P voltage imaging datasets in organotypic slices required slight deviations from this analysis pipeline. For data presented in Figure 2 and corresponding Supplement figure S5 each spatial filter was generated by regressing the whitened, detrended, segmentation masked, dataset against the corresponding electrophysiology traces which were taken as the ground-truth. Each electrophysiology trace was downsampled to match the acquisition rate of the voltage imaging data by slicing appropriate frequencies in the Fourier domain. For the data presented in Supplement figure S11, the binary segmentation mask was generated using the data acquired at 940 nm and was also applied to the corresponding data acquired at 1030 nm. All traces and results presented in Supplementary figure 1S1 were calculated using the unweighted average of pixels within the convex hull of the binary segmentation mask (the initial fluorescence trace described previously) and detrended using the adaptive b-spline approach.

For all imaging data sets that involved titration of the signal with the irradiance, we did not use the spatiotemporal correlation, but the same binary segmentation mask for all power densities. The details of this analysis are outlined in the respective methods section.

### Two-photon voltage imaging

The first tests for 2P voltage imaging with rhodopsin-based sensors in organotypic hippocampal slices were performed using temporally-focused holography as described before^12^ (Figure 2, Supplement figure S5). For further characterisation, we choose one of the speckle free methods like Generalised Phase Contrast (Supplement figure S6) or generated a low-NA Gaussian spot for the comparison of scanning and parallel illumination (Figure 3 and Supplement figure S7, S8, S11) and for the experiments in zebrafish larvae (Figure 4 and Supplement figure S9, S10) to obtain a homogeneous intensity profile of the spot. The details for each configuration are outlined below.

#### Holographic imaging in organotypic slices

Holographic scanless 2P voltage imaging was performed using temporally focused, holographic spots (lateral area: 335±16 μm^2^, axial extent: 9.6±1.6 μm) as described previously^12^. Light from an ultrafast source (Coherent Discovery, ∼1 W, 80 MHz, 100 fs tuned to 940 nm) passed through a half-wave plate (Thorlabs, WPHSM05-980) mounted on a motorised rotation mount (Thorlabs PRM1Z8) and polarising beam splitter (Thorlabs, CCM1-PBS253/M). The orientation of the half-wave plate was controlled during experiments to modulate the average power. The beam was expanded using a Galilean beam expander formed of two lenses (focal length = −75 mm, Thorlabs, LC1258-B and focal length = 500 mm Thorlabs, AC508-500-B), resulting in a beam which slightly underfilled the SLM screen (Hamamatsu, LCOS X13138-0, 1272 × 1024 pixels, 12.5 μm pitch). Holograms were calculated using an iterative Gerchberg-Saxton algorithm. The modulated beam was Fourier transformed using a lens with focal length 750 mm (Thorlabs, AC508-750-BG2) onto a blazed diffraction grating (600 lines/mm, Thorlabs GR50-0610L11). The illumination angle of the diffraction grating was chosen in order that the first diffraction order of the central wavelength (940 nm) propagated along the optical axis such that the temporal focusing plane coincided with the imaging plane of the microscope. Finally, a pair of telescopes was used to de-magnify and relay the holographic spots onto the focal plane (focal length = 500 mm Thorlabs, AC508-500-BL12 Lens, focal length = 300 mm, Thorlabs, AC508-300-B, Objective lens, 40X, 0.8 NA, f = 5 mm, Nikon, CFI APO NI). The zeroth diffraction order was removed using a physical beam block positioned in the conjugate image plane following the grating. Fluorescence was captured using a simple wide field detection axis comprised of the same microscope objective, a tube lens (Thorlabs, TTL200-A) and a scientific complementary metal-oxide semiconductor (sCMOS) camera (Photometrics Kinetix) controlled using micro-manager 2.0. Fluorescence was filtered using either a quad-band filter (Chroma ZET405/488/561/640) or individual bandpass filters. Infrared light was blocked from the camera using a shortpass filter.

Imaging data was analysed as described in “Analysis of imaging data” above.

#### Two-photon scanning *vs*. parallel illumination in cultured cells

Recordings were performed at 940 nm using a Ti:Sapphire Laser (Mai-Tai Spectra Physics, 80MHz repetition rate) and a customised microscope providing multiple illumination paths (Supplement figure S7A). In a first scanning path, after Pockels cell modulation (Conoptics), the beam was directed toward a commercial galvo-galvo scanning head (VIVO 2-PHOTON operated by Slidebook 6 software, 3i), to achieve a fast raster scan over the cell body. After a first relay lens (f_L3_=5 cm and f_L4_=25 cm) and the microscope objective (W Plan-Apochromat 20×, NA 1.0, Zeiss), the beam was focused on the sample with a lateral 2P-PSF size of ≈ 0.9 μm (1/e^2^), which was used to calculate the illuminated surface to be approximately 0.64 μm^2^. Excitation power ranged between 15 and 70 mW out of the objective corresponding to irradiances from 23 to 109 mW/μm^2^.

In the parallel illumination path, after a first beam reducer (f_L1_ =50 cm and f_L2_ =20 cm), a collimated Gaussian beam was diaphragmed by an iris, which plane was demagnified and conjugated with the sample plane (using L_4_ and the objective mentioned above) to project a top hat circular spot with a radius of ≈ 15 μm (729±15 μm^2^), resulting in an irradiance of 0.4 ±0.01 mW/μm^2^.

The parallel spot and the scan trajectories were matched to illuminate a similar portion of the cell body (Supplement figure S7B). The two paths were recombined using a polarising beam splitter and the beam was alternatively directed towards one of them by a motorised flip mirror. In both cases, fluorescent emission was detected by a PMT (band pass filter: 510/84, Semrock) (Supplement figure S11). Additionally, camera detection with an CMOS camera was available, together with patch clamp recording setup as described above.

Resulting data was exported to tiff-format and further analysed using custom pythons scripts. For data acquired with the PMTs (Figure 3 and Supplement figure S11), resulting kymographs were averaged along the spatial axis without excluding any lines to obtain the average fluorescence trace (Supplement figure S7). F_0_ was determined as the average fluorescence over 55 timepoints after ∼200 ms of illumination (V_hold_ = −70 mV) and the standard deviation SD_F(−70 mV)_ was determined as standard deviation of the fluorescence over the same range. The relative fluorescence change was calculated as 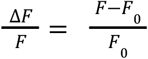 while the z-score was calculated as 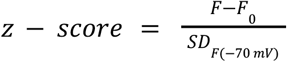. The resulting traces were neither detrended nor corrected for bleaching unless indicated otherwise, but for the 100 ms depolarisations before extraction of the average ΔF/F_0_ and z-score per voltage step, a Gaussian filter (sigma = 3) was applied to the trace. The displayed traces are unfiltered.

For data acquired with the CMOS camera (DaVinci2K) the cell was manually segmented based on the image stack acquired with parallel illumination and the fluorescence was averaged over the segmented region of interest (Supplement figure S8). The same ROI was used for recordings with scanning illumination on the same cell. A region outside of the ROI was used to calculate the background and correct the fluorescence trace. The fluorescence traces were then processed as described above. Here, traces for display were filtered with a 1^st^ order butterworth lowpass filter.

#### Generalised Phase Contrast imaging in cultured cells

Patch-clamp experiments with illumination using GPC (Supplement figure S6) were performed in ND7/23 cells expressing Jarvis, pAce or JEDI as described above. The setup and realisation of the GPC spot are described in Sims and Bendifallah *et al*.^12^. Here, we projected a large (305 ± 5 μm^2^) temporally-focused spot with an axial resolution of 4.3±0.1 μm using a Coherent Chameleon Discovery laser (1.2 W, 80 MHz repetition rate) tuned to 940 nm. Powers were measured out of the objective and power densities ranging from 0.2 to 1.0 mW/μm^2^ calculated accordingly. Data was acquired with an sCMOS camera (Kinetix, Photometrics) at frame rates of 50 or 1000 Hz as indicated.

Resulting data was analysed similar to the camera data acquired for the comparison of scanning and parallel illumination (see above) with the difference that a linear detrending around the voltage step was applied here to account for bleaching.

#### Illumination with a low-NA Gaussian spot in zebrafish larvae

4-6 dpf zebrafish larvae transiently expressing Jarvis were paralyzed by 3-5 minutes incubation in α-bungarotoxin (1 mg/mL) and mounted in a petri dish using 2% (v/v) low melting point agarose. Jarvis-expressing, spontaneously spiking olfactory sensory neurons were identified by 1P wide field imaging (described above; 470 nm LED, 7 mW/mm^2^) with an sCMOS camera (Kinetix, Photometrics) at a frame rate of 991 Hz. Active neurons were subsequently imaged under parallel 2P illumination with a temporally-focused low-NA Gaussian spot (137±10 μm^2^) using a Ti:Sapphire laser (MaiTai, Spectra Physics) tuned to 920 nm. Temporal focusing was achieved using a blazed diffraction grating with 830 lines/mm (Optometrics, Littleton, MA, USA) resulting in an axial extent of the spot of 8 μm (FWHM). The average power out of the objective was 80-100 mW corresponding to power densities between 0.61-0.76 mW/μm^2^.

Data analysis was performed as described in “Analysis of imaging data”.

### Statistics and figures

If not indicated otherwise, all data is represented as mean±standard deviation (SD) together with individual recordings. For direct comparisons (expressing *vs*. non-expressing or cells expressing different sensors) few recordings for all groups were performed on the same recording day, on the same batch of samples. For all *in-vitro* experiments, data was collected from at least two independent transfections or transductions on at least two different days. Statistical analysis was performed using the Mann-Whitney-U or Wilcoxon rank-sum test as indicated using custom python scripts (scipy package version 1.11.1).

Figures were prepared using Origin 2023 (version 10.0.0.154) and Affinity Designer (version 1.10.6).

## Acknowledgments

We thank Christoph Tourain for relentless engineering support, Vincent de Sars for IT support and Eric Schreiter, Ahmed Abdelfattah, Vincent Pieribone, and Emiliano Ronzitti for fruitful discussions. We are further very grateful for the support of Peter Hegemann with cloning and the encouraging words of Lawrence B. Cohen in the early stages of the project. We thank Marnie Halper for sharing the plasmid containing the original *Xla*.*Tubb2* promoter sequence.

We acknowledge the contributions of the viral vector facility at the Institut de la Vision, the work of the animal facility and the support of Stéphane Fouquet with confocal microscopy.

This work was supported by the Agence National de Recherche (ANR-19-CE16-0026, ‘HOLOPTOGEN’) (VE), and the European Research Council (ERC2019-ADG-885090, ‘HOLOVIS-AdG’) (VE); the European Union’s Horizon 2020 programme under the Marie Skłodowska-Curie grant 813457 (CC and EleP), Fondation pour la Recherche Médicale FDT-Fin de thèse FDT202304016818 (CC), Fondation de France, Maladies de l’oeil (00111014) (GF); CG was supported by the Deutsche Forschungsgemeinschaft DFG (442616457) during parts of this project.

## Author Contributions

CG and VE conceived the project. CG designed and cloned the sensor variants and performed the one-photon *in-vitro* comparison with the help of ASM. The setup used for two-photon holographic imaging was built and maintained by EirP and RRS, the organotypic slices were prepared by ASM, the patch-clamp/imaging experiments were performed by CG and corresponding data analysis was done by RRS. Confocal imaging was performed by ASM. The setup for comparing scanning and parallel illumination was built and maintained by DT, with experiment design and interpretation by DT and CG, experiments and data analysis were performed by CG supported by DT and RRS. GF subcloned the sensor for zebrafish expression and injected the line generated by ElP, under guidance of FDB; the setup for zebrafish experiments was implemented by CC corresponding imaging was performed by CC supported by CG while data was analysed by RRS. The manuscript was written by CG and VE with contributions from all authors. VE raised funding and supervised the project.

## Declaration of interests

The authors declare no competing interests.

## Notes

### Competing Interest Statement

The authors have declared no competing interest.

