## Supplement Material for "Two-photon voltage imaging with rhodopsin-based sensors"

Supplement figures S1- S12

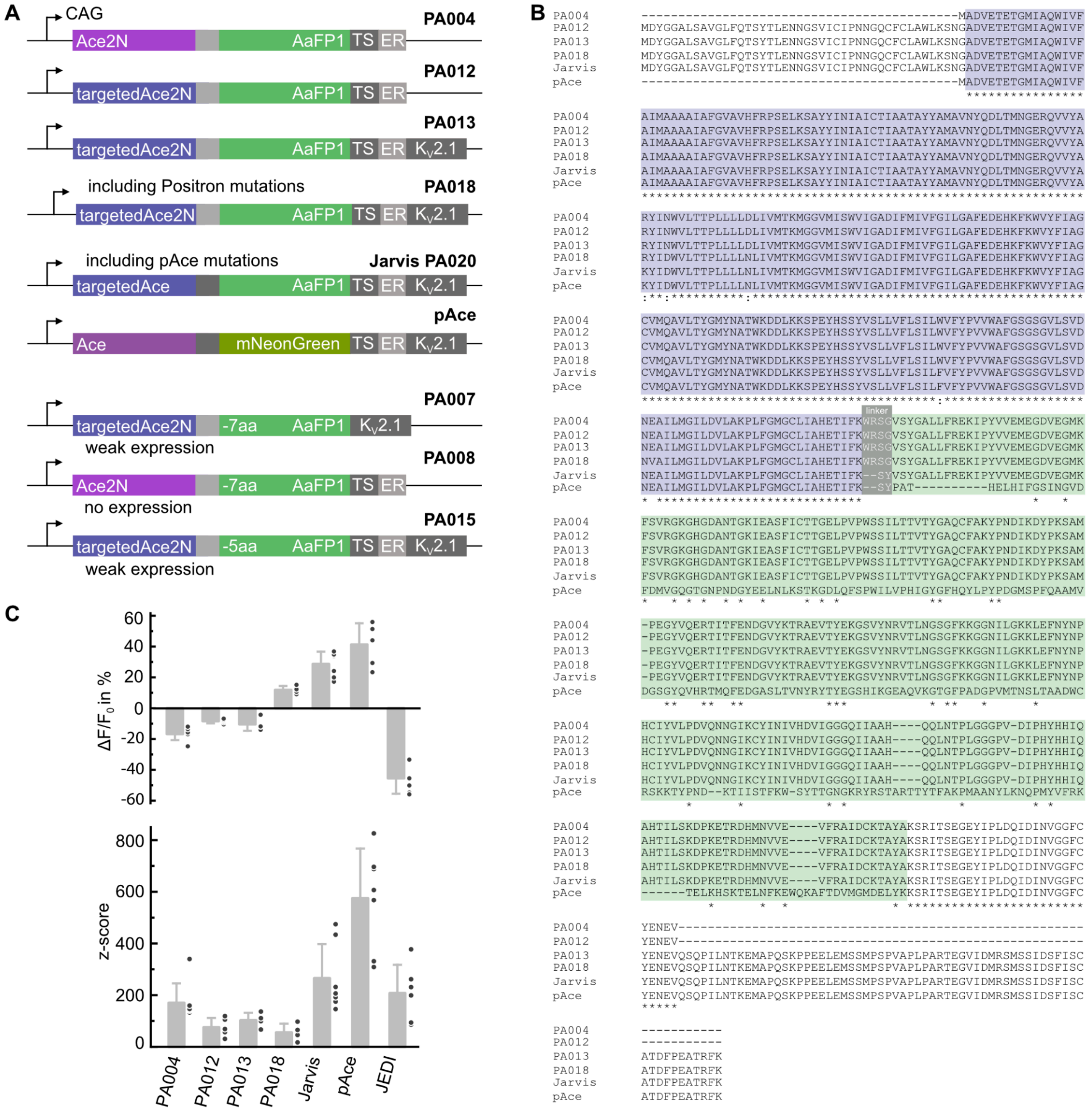


**Supplement figure S1: Rational engineering of Jarvis A)** Molecular design and **B)** alignment of DNA sequences of selected variants of Ace (purple) and AaFP1 (green) fusions, Jarvis and pAce. **C)** *In-vitro* characterisation of select variants compared to Jarvis, pAce, and JEDI-2P in cultured ND7/23 cells using 1P widefield illumination (470 nm LED, 10 mW/mm$^2$) with camera detection (50 Hz) and simultaneous whole-cell voltage-clamp (-70 mV to 30 mV for 100 ms). $\Delta F/F_0$ and z-score extracted from imaging traces in the middle of the voltage step. Points show individual recordings and bars represent mean±SD.

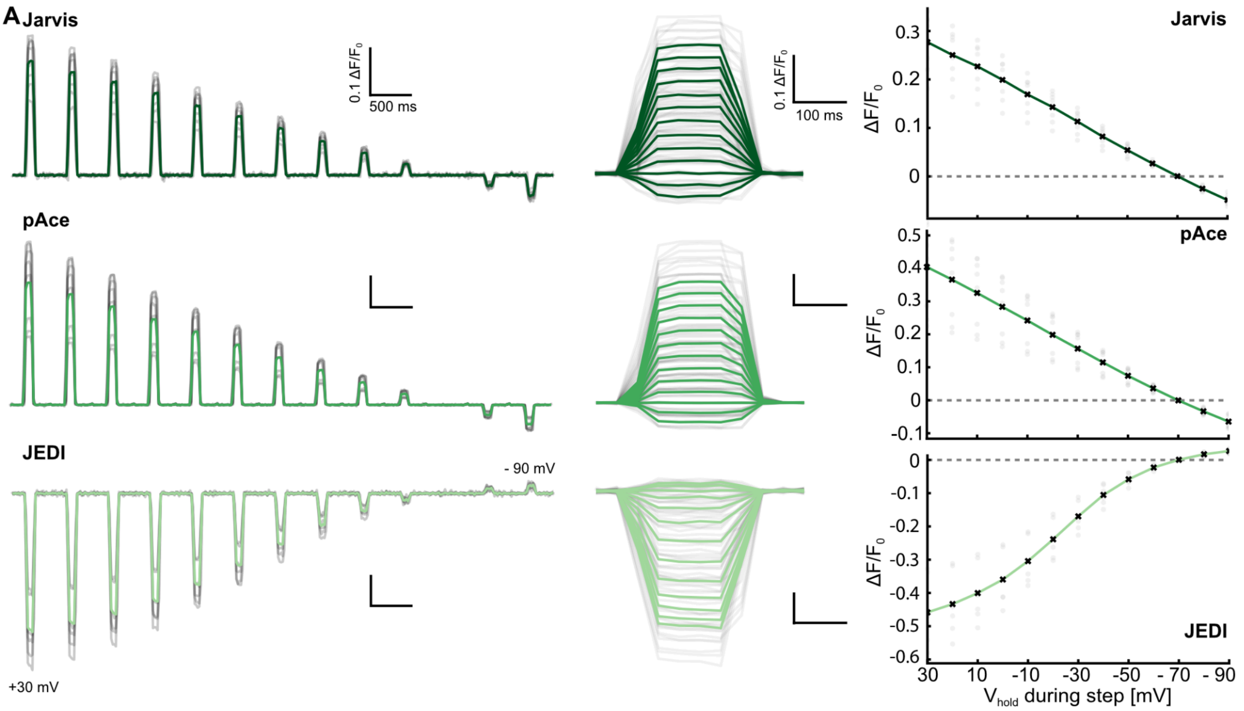


**Supplement figure S2**: 1P sensitivity curves of Jarvis, pAce, and JEDI-2P in cultured ND7/23 cells using 1P widefield illumination (470 nm LED, 10 mW/mm$^2$) with camera detection (50 Hz) and simultaneous whole-cell voltage-clamp to step the holding potential from -70 mV (100 ms) in the range from + 30 mV to -90 mV. Average $\Delta F/F_0$ trace (green) with all individual recordings (grey).

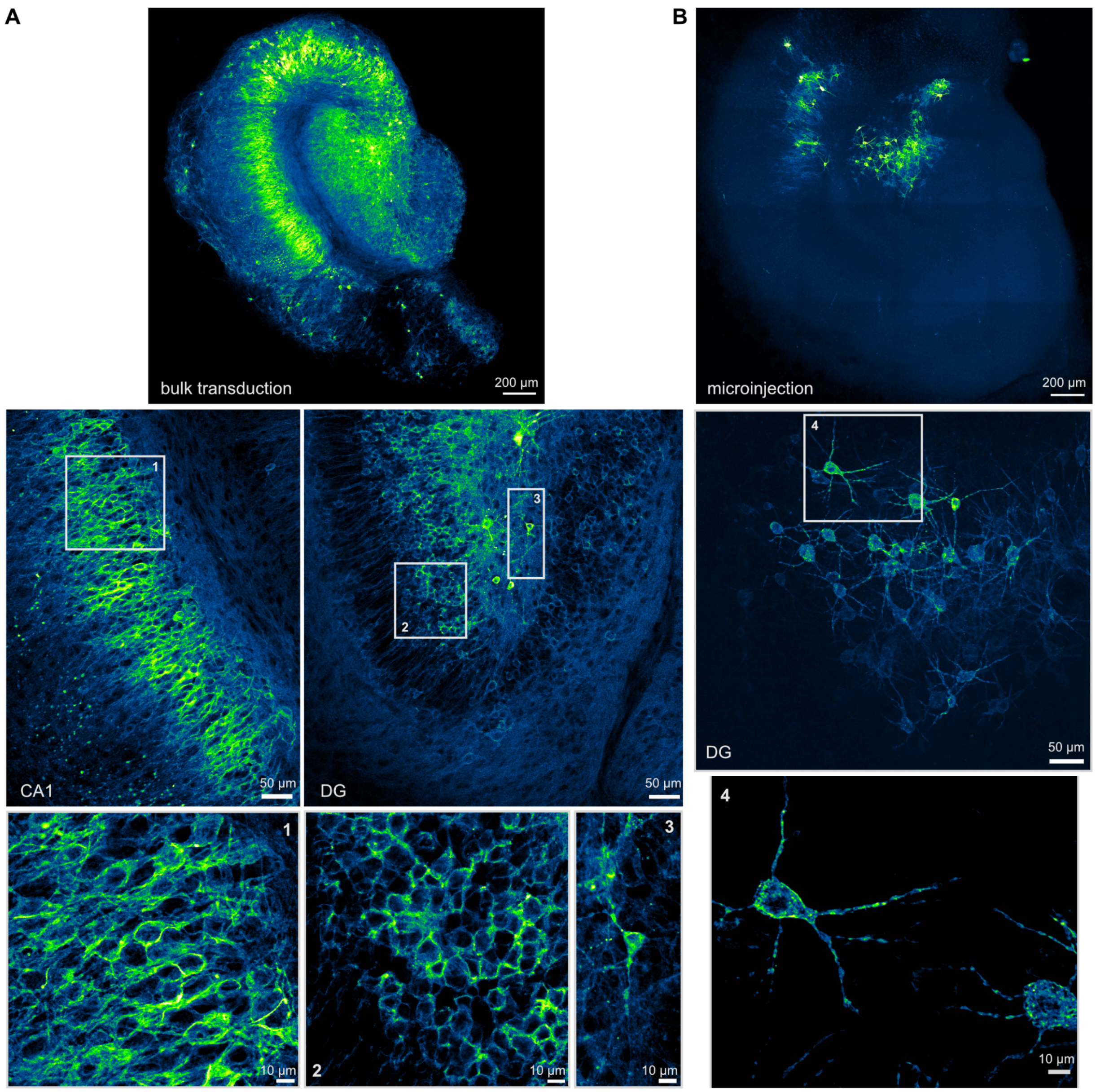


**Supplement figure S3: Confocal images of fixed organotypic hippocampal slices expressing Jarvis after viral transduction. A** Top: Confocal image of a whole bulk-transduced slice expressing Jarvis in all regions; maximum projection of 37 planes (0.6 µm each). Zoom-in at higher resolution for CA1 and dentate gyrus (middle) and for a few cells in each region (bottom); average projections of 3-4 planes of 2 µm thickness. **B** Top: Confocal image of a whole microinjected slice expressing Jarvis localised around the microinjection site in DG and a small part of CA1; maximum projection of 35 planes (0.6 µm each). Zoom-in at higher resolution for the microinjected region (middle) and two cells within that (bottom); average projections of 3-4 planes of 2 µm thickness.

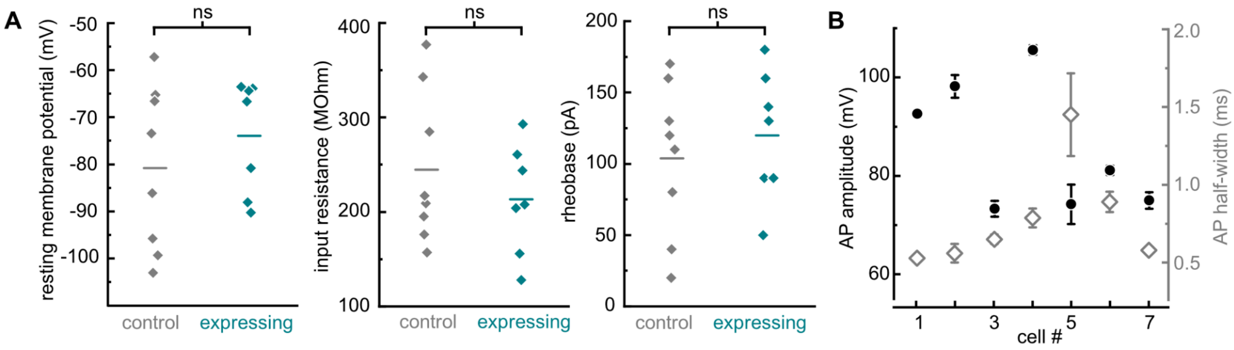


**Supplement figure S4: Electrical properties of Jarvis-expressing neurons in organotypic slices after viral transduction.** Bath solution heated to 33 °C and with 4 mM $CaCl_2$ and 4 mM $MgCl_2$ to reduce excitability of the slices. **A)** Membrane properties of Jarvis-expressing neurons (teal) compared to non-expressing neurons (grey) in the same preparation. Left: Resting membrane potential immediately after going whole-cell; LJP corrected. Middle: Input resistance as slope of the linear fit of the voltage-current relation for current injections from -30 pA to +30 pA in 10 pA steps. Right: Rheobase as the lowest 1 s long square current-injection that evoked an AP. All data points are displayed and the line indicates the average value. No significant differences (Mann-Whitney U test) were found for any of the parameters between Jarvis-expressing and non-expressing neurons. **B)** Amplitude (black) and half-width (grey) of APs extracted from the electrophysiology traces corresponding to the optical recordings (1000 Hz acquisition) presented in figure 2. Displayed as mean±SD with 9-13 APs per neuron.

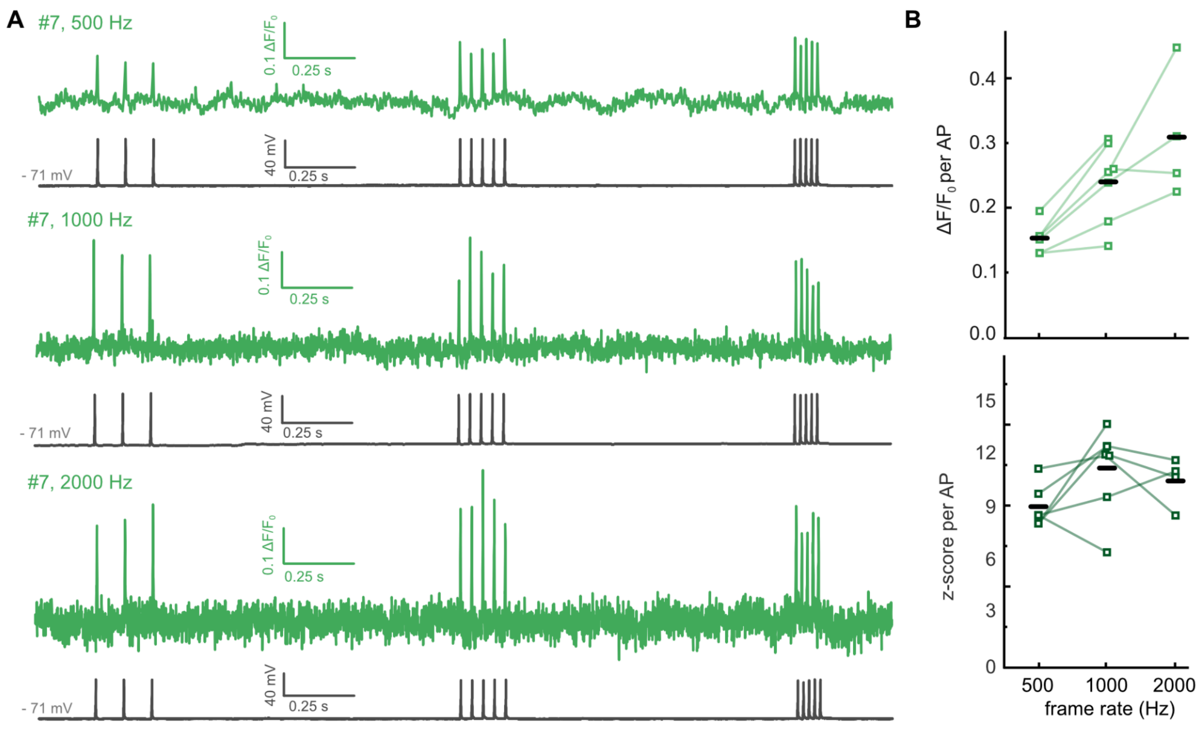


**Supplement figure S5: AP detection with Jarvis at different frame rates.** Illumination as in Figure 2 (temporally-focused holographic spot, 940 nm, ~20 µm diameter, ~10 µm axial extent, 0.66 mW/µm$^2$) **A)** $\Delta F/F_0$ (green) and corresponding electrophysiology traces (grey) for the same neuron (#7) imaged at 500 Hz, 1000 Hz and 2000 Hz. **B) (i)** Summary of $\Delta F/F_0$ and z-score per AP at different frame rates for the same neuron, respectively. Squares show the average from 8-13 APs per neuron; black lines indicate the average over all neurons for the respective frame rate.

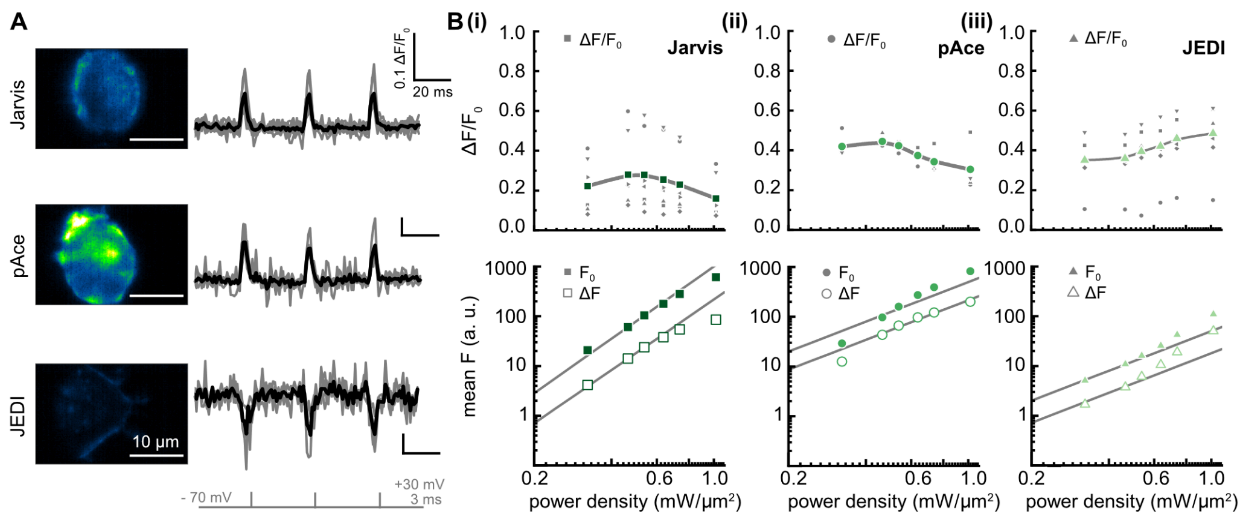


**Supplement figure S6: Two-photon imaging with Generalised Phase Contrast.** ND7/23 cells expressing Jarvis, pAce or JEDI were patched (whole-cell configuration) and the holding potential was stepped from - 70 mV to +30 mV **A)** for 3 ms (three times) while cells were illuminated with a large temporally-focused GPC spot (940 nm, ~18 µm diameter, ~5 µm axial extent, 1.03 mW/µm$^2$) and imaged with the Kinetix camera at 1 kHz. Grey traces show individual recordings while black traces show the average $\Delta F/F_0$ trace; n = 4-7 each construct. **B)** $\Delta F/F_0$ (top) and average F (bottom) at rising power densities from 0.33 mW/µm$^2$ to 1.03 mW/µm$^2$ resulting from 100 ms depolarisations (-70 mV to +30 mV) for (i) Jarvis (n = 7), (ii) pAce (n = 4) and (iii) JEDI (n = 7) expressing cells; acquisition at 50 Hz. Top panels show average values (coloured) and individual data points (grey). Lower panels show average values for $F_0$ and absolute fluorescence change $\Delta F$; lines have a fixed slope of 2.

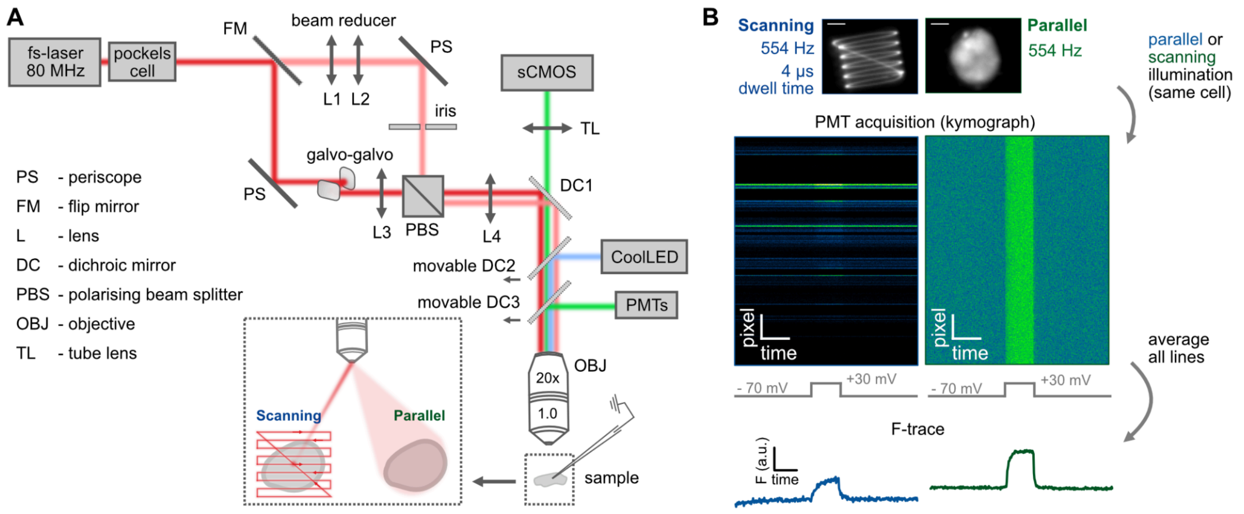


**Supplement figure S7: Setup for comparison of scanning and parallel illumination. A)** Microscope design allowing for parallel or scanning illumination over the same region. For scanning illumination we defined a square-shaped trajectory that was scanned with a diffraction limited spot at a speed of 1.84 ms per line (4 µs per location) corresponding to 554 Hz. For the parallel illumination we created a low-NA Gaussian spot over the same region with an area of 730 $\mu m^2$. In both cases acquisition was performed with PMTs. **B)** Top: Resulting scanning or parallel illumination visualised by exciting a thin rhodamine layer and taking an image with a camera (1.84 ms exposure); both images show the same field of view, scale bar 10 µm. Bottom: Representative kymographs acquired during a recording with scanning (left) and parallel (right) illumination. The cells were patched and the membrane voltage was stepped from -70 mV to +30 mV for 100 ms. For analysis kymographs were averaged along the lines to get the average fluorescence trace; no lines excluded.

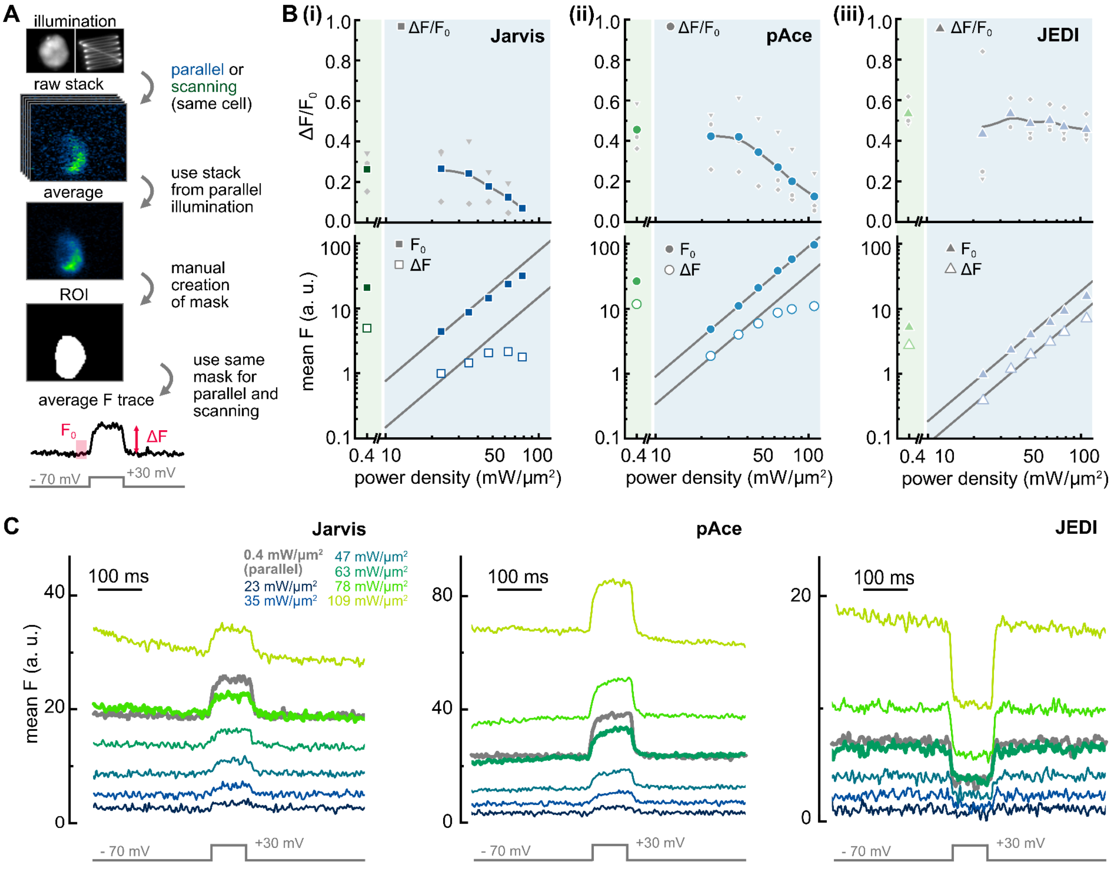


**Supplement figure S8: Comparison of scanning and parallel illumination with camera detection.** Illumination, samples and recordings are set up as in Figure 3 with the difference that a CMOS camera was used for detection; power densities and frame rate (554 Hz) are the same as in Figure 3. **A)** Schematic of recording protocol and data analysis pipeline. ND7/23 cells are patched and the holding voltage is stepped from - 70 mV to 30 mV for 100 ms while the voltage change is imaged simultaneously. From the image stack received with parallel illumination the cell is segmented manually to create the mask, which is then also used for the recordings with scanning illumination. Fluorescence is then averaged over the mask to generate the fluorescence trace, as described in detail in materials and methods. **B)** $\Delta F/F_0$ (top) and average F (bottom) at rising scanning powers from 23 mW/µm$^2$ to 109 mW/µm$^2$ (blue) and corresponding value from parallel illumination (green) for (i) Jarvis, (ii) pAce and (iii) JEDI; n = 3 each. For $\Delta F/F_0$ (top) the line does not represent a fit, just guidance to the eye. For the average F it shows a line with a fixed slope of 2. **C)** Raw imaging traces (no detrending, no denoising) acquired for recordings presented in B) for Jarvis, pAce and JEDI with the F-trace from the parallel illumination in grey and rising scanning powers as indicated. Highlighted (bold) are the parallel F-trace and the scanning power that resulted in comparable baseline fluorescence $F_0$.

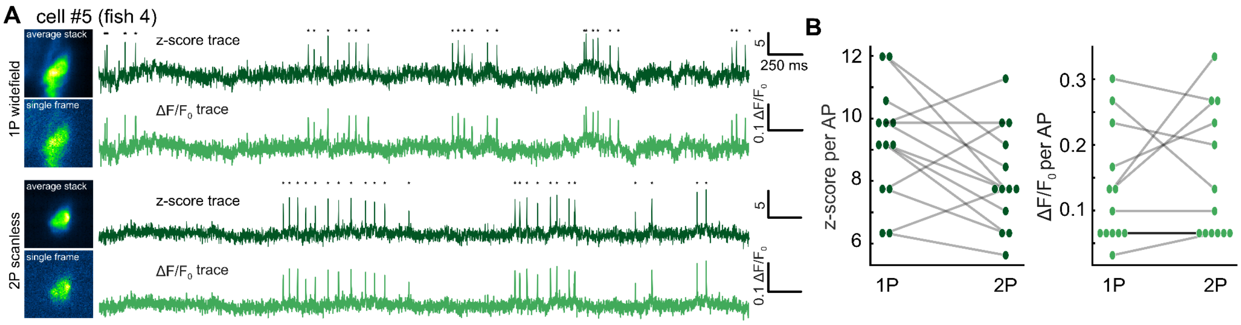


**Supplement Figure S9: 1P widefield and 2P scanless illumination for AP detection with Jarvis in zebrafish larvae. A)** Exemplary recordings with resulting z-score (dark green) and $\Delta F/F_0$ traces (light green) for the same neuron under 1P widefield (top) and 2P scanless (bottom) illumination. **B)** Average z-score and $\Delta F/F_0$ per AP under 1P and 2P illumination. Data for the same 13 neurons that are presented in Figure 4D.

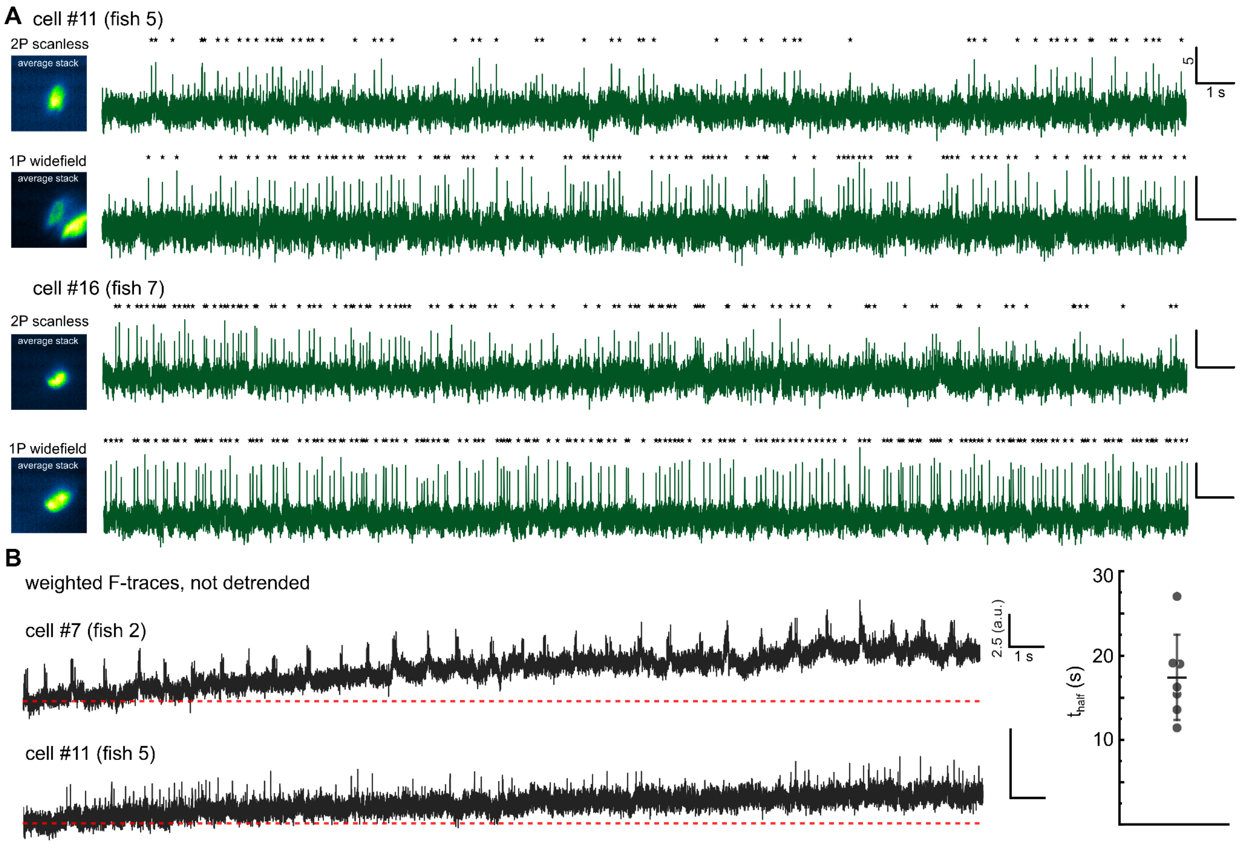


**Supplement Figure S10: AP detection with Jarvis in zebrafish larvae over 30 s. A)** Exemplary z-score traces for two cells resulting from 1P widefield (bottom, 470 nm, 7 mW/mm$^2$) and 2P scanless illumination (top, 920 nm, 0.61- 0.76 mW/µm$^2$); acquired with an sCMOS camera at 991 Hz over 30 s. **B)** Non-detrended weighted F-traces for two cells exemplifying a slow rise of $F_0$ (photoactivation) during the recording with 2P scanless illumination and an estimation of the half life time for several cells (right).

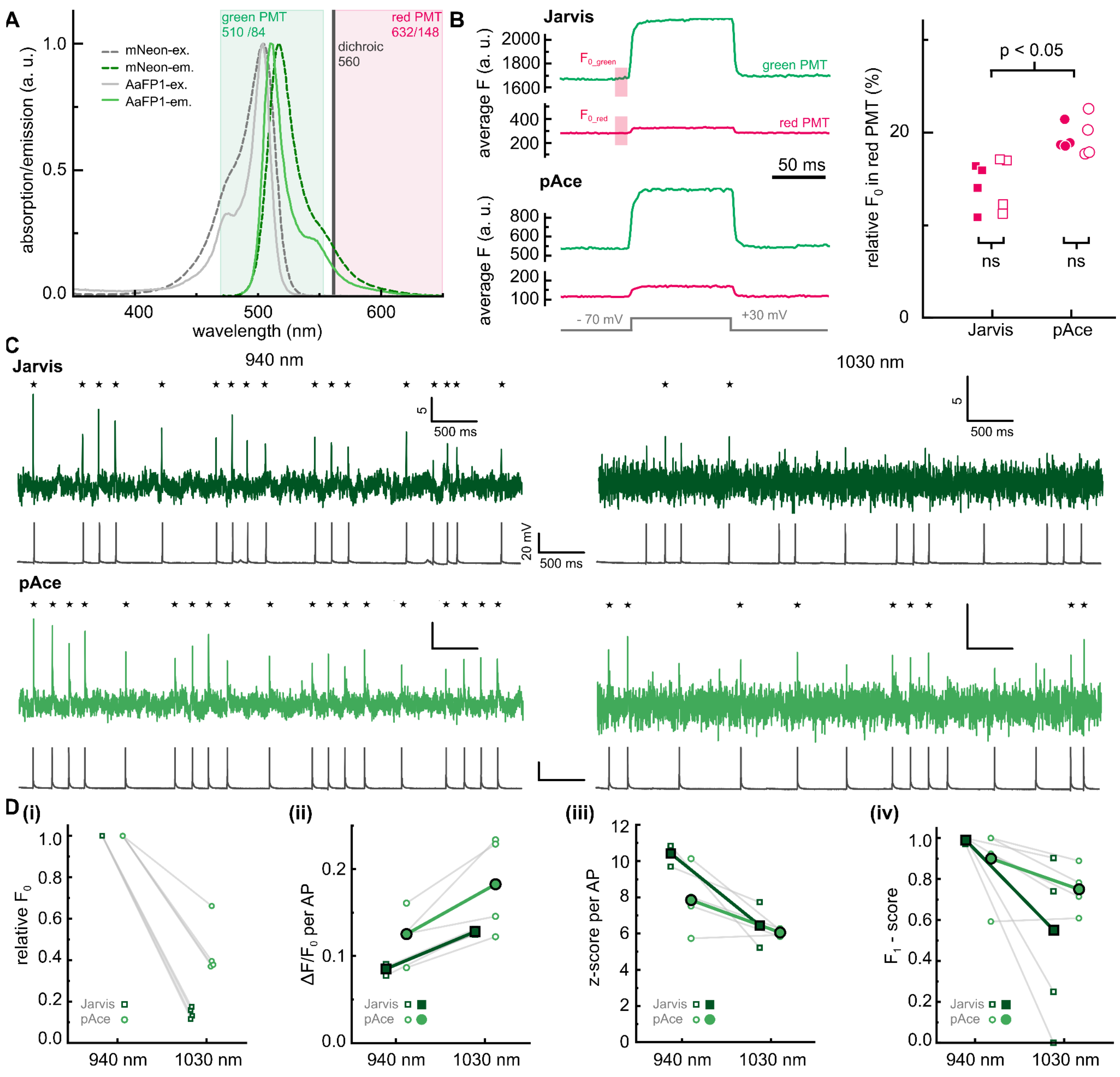


**Supplement figure S11: Two colour imaging with Jarvis *vs.* pAce. A)** Excitation and emission spectra of *Aa*FP1 and mNeonGreen (re-plotted from FPbase.org), bandwidth of emission filters for the PMTs and dichroic mirror. **B)** Fluorescence traces for Jarvis (top) and pAce (bottom) acquiring on both PMTs. **C)** Average $F_0$ on the red relative to the green PMT ($F_{0,red}/F_{0,green}$) for Jarvis (squares) and pAce (circles). Four cells each; open symbols scanning and filled symbols parallel illumination; no significant difference between the illumination modes. There was a significant difference between Jarvis 14±3% and pAce 20±2% (Wilcoxon rank-sum test). **C)** Z-score traces (green) from scanless voltage imaging in organotypic hippocampal slices expressing Jarvis (dark green) or pAce (light green) after viral transduction using a non-temporally focused holographic spot at 940 nm (left) or 1030 nm (right) and camera detection at 1 kHz; simultaneous current-clamp recordings (grey) to monitor membrane potential and evoke APs. Powers adjusted to equal photon flux at both wavelengths and same hologram was used resulting in spot diameters and power densities of 14 µm, 0.8 mW/µm$^2$ at 940 nm and 15.5 µm, 0.6 mW/µm$^2$ at 1030 nm. Both wavelengths recorded on the same cell in randomised order across cells. **D)** AP detection (*) using only the optical trace (see methods section for details); same segmentation for both wavelengths. (i) Relative $F_0$ averaged over the first 50 ms of illumination. **(ii)** $\Delta F/F_0$ and **(iii)** z-score per detected AP. **(iv)** Detectability of APs in the optical trace as $F_1$-score calculated according to $F_1$ = 2*TP/(2*TP+FN+FP) with TP-true positive, FN-false negative and FP-false positive (verified with electrophysiology).

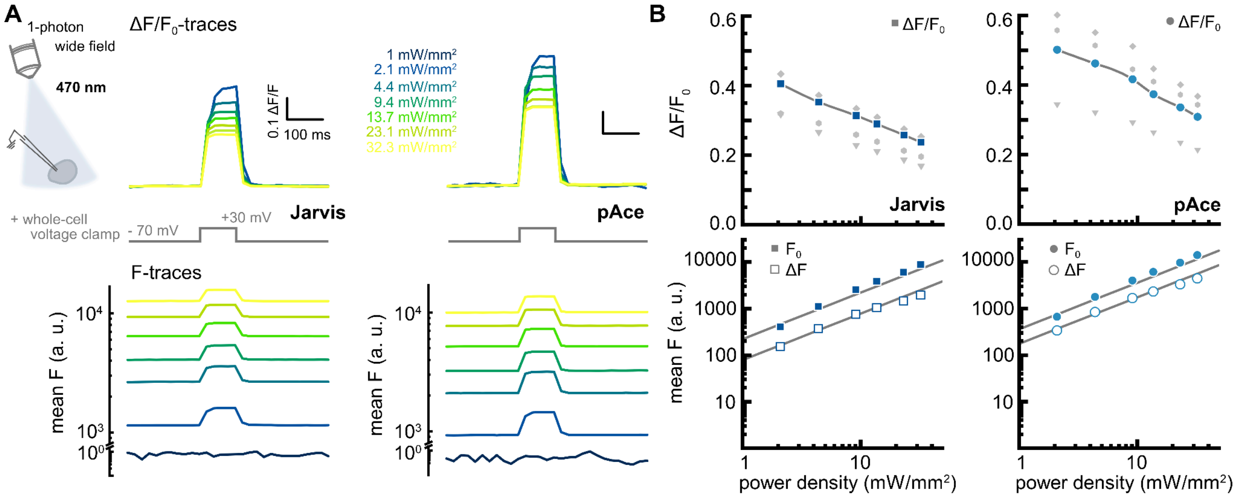


**Supplement figure S12: Titration of $\Delta F/F_0$ for a 100 mV step with the 1P widefield irradiance. A)** Whole-cell voltage clamp recordings were performed on cultured ND7/23 cells expressing Jarvis (left) or pAce (right) to step the membrane voltage from -70 mV to +30 mV and optical traces were acquired simultaneously with an sCMOS camera (50 Hz) using 1P wide field illumination of rising irradiances (470 nm LED). Displayed are two exemplary traces of $\Delta F/F_0$ (top) and for the average fluorescence F (bottom) of one cell at different irradiances. Cell were segmented based on fluorescence and the same segmentation was used for all power densities; there was no detrending, denoising or pixel weighting in this analysis. **B) Top:** $\Delta F/F_0$ in dependence of the used power density (top) for Jarvis (left) and pAce (right); coloured symbols are the average of three cells while grey ones denote individual data points. The line is not a fit but guidance to the eye. **Bottom:** Raw average fluorescence values (filled) together with the respective absolute fluorescence change ΔF (open). Grey lines have a fixed slope of 1 to indicate the expected signal rise for 1P excitation.
